# Gut microbiome-based prediction of autoimmune neuroinflammation

**DOI:** 10.1101/2023.04.14.536901

**Authors:** Alex Steimle, Mareike Neumann, Erica T. Grant, Stéphanie Willieme, Alessandro De Sciscio, Amy Parrish, Markus Ollert, Eiji Miyauchi, Tomoyoshi Soga, Shinji Fukuda, Hiroshi Ohno, Mahesh S. Desai

## Abstract

Gut commensals are linked to neurodegenerative diseases, yet little is known about causal and functional roles of microbial risk factors in the gut–brain axis. Here, we employed a pre-clinical model of multiple sclerosis in mice harboring distinct complex microbiotas and six defined strain combinations of a functionally-characterized synthetic human microbiota. Discrete microbiota compositions resulted in different probabilities for development of severe autoimmune neuroinflammation. Nevertheless, assessing presence or the relative abundances of a suspected microbial risk factor failed to predict disease courses across different microbiota compositions. Importantly, we found considerable inter-individual disease course variations between mice harboring the same microbiota. Evaluation of multiple microbiome-associated functional characteristics and host immune responses demonstrated that the immunoglobulin A-coating index of *Bacteroides ovatus* before disease onset is a robust individual predictor for disease development. Our study highlights that the “microbial risk factor” concept needs to be seen in the context of a given microbial community network, and host-specific responses to that community must be considered when aiming for predicting disease risk based on microbiota characteristics.

## INTRODUCTION

Compared to healthy controls, autoimmune disease patients exhibit distinct microbiota compositions (*1*), especially in the context of multiple sclerosis (MS) (*2*). Thus, determining whether susceptibility or progression of MS can be predicted by the microbiota composition is a necessary precondition to develop patient-targeted microbiota modulations (**Fig. 1a**). A common approach to elucidate microbiota-related, MS-promoting predictors compares relative abundances of bacterial taxa – often determined by 16S rRNA gene-based sequencing – between MS-affected and healthy individuals (*2–9*). Although certain differentially abundant taxa identified across different human cohort studies tend to be concordant, e.g., increased abundances of *Akkermansia* (*2, 4, 5, 7–9*) or decreased abundances of *Prevotella* (*3, 5–8*) in MS patients compared to healthy controls, such cohort-level observations do not explain inter-individual differences (*10*) in disease course or susceptibility. Therefore, it remains challenging to reliably link taxa abundances across individuals to microbiota characteristics that impact MS disease course.

**Figure 1:**
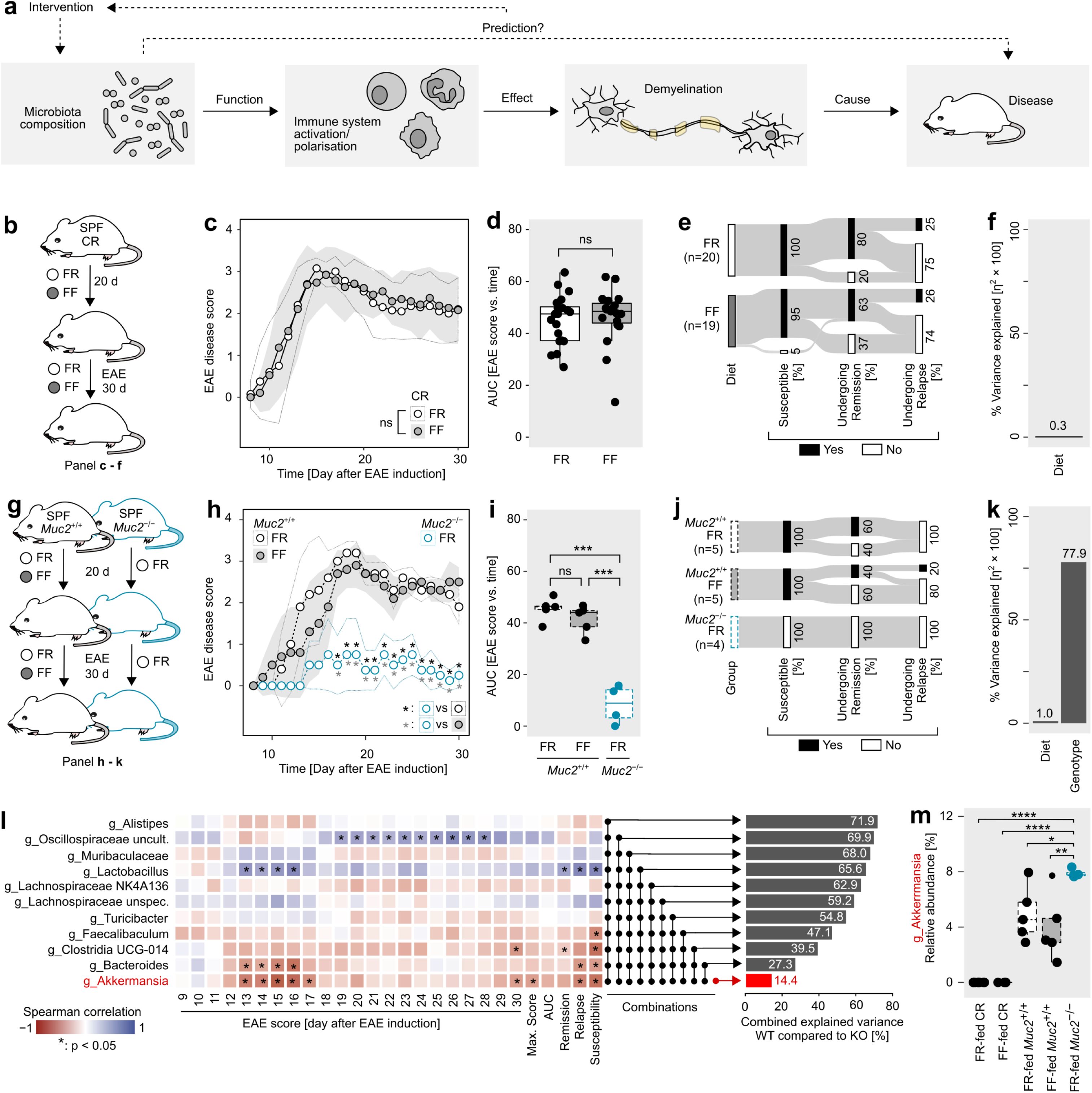
Increased levels of *Akkermansia* are associated with lower neuroinflammation in EAE-induced mice harboring a complex native microbiota. **a)** Central objective of the study: To investigate how microbiota composition interacts with the host immune system (exhibits a specific “function”), influences the degree of demyelination during experimental autoimmune encephalomyelitis (EAE) and results in different disease phenotypes. Microbiota composition-based prediction of disease-mediating properties would make targeted microbiota manipulation (“intervention”) possible. **b)** Experimental setting for panels c) to f). C57BL/6J mice purchased from Charles River (CR), housed under SPF conditions, were fed either a fiber-rich (FR) or fiber-free (FF) diet for 20 d, followed by induction of EAE. Mice were kept on the respective diets during EAE course. EAE disease development was recorded for 30 d. **c)** EAE disease scores as a function of time. All mice (FR, n=20; FF, n=19; two independent experiments) were scored daily at the same time. Dots represent group means; shaded area and dotted lines represent SD. Statistics: daily EAE scores were compared using a Wilcoxon rank-sum test. “ns”: no statistically significant difference between groups at any day. **d)** Area-under-the-curve (AUC) analysis of the disease course depicted in c). Each mouse depicted by a separate dot. Statistics: unpaired t-test after verification of normal distribution of values using a Shapiro-Wilk test. Boxplots follow standard Tukey representations. **e)** Sankey diagram of key event occurrence (in % of all mice within one group) during EAE. Susceptibility: score of 2.5 for at least 1 d. Remission: decrease of EAE score by 1.5 points compared to maximum score. Relapse: Increase by 1.0 point compared to remission score. **f)** Percentage of variance explained by diet when comparing AUC between FR and FF groups. Determined by eta-squared (η^2^) calculation. **g)** Experimental setting for panels h) to k). *Muc2*^+/+^ and *Muc2*^−/−^ littermate mice were fed an FR (*Muc2*^+/+^ and *Muc2*^−/−^) or FF (*Muc2*^+/+^ only) diet for 20 d, followed by EAE induction and observation of disease course for 30 d. Mice were kept on their respective diets during EAE. **h)** EAE disease scores as a function of time. All mice (FR-fed *Muc2*^+/+^, n=5; FF-fed *Muc2*^+/+^, n=5; FR-fed *Muc2*^−/−^, n=4) were scored daily at the same time. Dots represent group means; shaded area and dotted lines represent SD. Statistics: daily EAE scores compared using a Wilcoxon rank-sum test followed by Benjamini-Hochberg p-value correction for multiple comparisons. *:p<0.05. Grey asterisks: FR-fed *Muc2*^−/−^ vs FR-fed *Muc2*^+/+^, Black asterisks: FR-fed *Muc2*^−/−^ vs FF-fed *Muc2*^+/+^. **i)** AUC analysis of the disease course depicted in h). Each mouse depicted by a separate point. Statistics: One-way ANOVA followed by Tukey’s post-hoc test for groupwise comparison. ns: p > 0.05; ***, p < 0.001. **j)** Sankey diagram of key event occurrence (in % of all mice within one group) during EAE. Same event definitions as in e). **k)** Percentage of variance explained by diet and genotype when comparing AUC between the three groups depicted in h), as determined by eta-squared (η^2^) calculation. **l+m)** Combined analysis of microbial communities of CR mice (FF- and FR-fed), *Muc2*^+/+^ mice (FF- and FR-fed) and FR-fed *Muc2*^−/−^ mice as determined by Illumina-based 16S rRNA gene sequencing from fecal samples collected before EAE induction. All samples were analyzed together using the same analysis pipeline. **l)** Genera, which in summary explained more than 70% of the variance, as calculated based on Bray-Curtis dissimilarity index obtained from relative abundance data on a genus level, of fecal microbial communities between WT (CR and *Muc2*^+/+^, fed both diets, combined) and KO (*Muc2*^−/−^) mice before EAE induction. Squares: Spearman correlations between relative abundances of the selected genera before EAE induction with EAE-associated readouts as calculated by pairing values within each individual mouse across all five groups. Statistical significant correlations indicated by *. Barplots depict cumulative explained variance for combinations of genera, ordered from bottom (highest single contribution) to top (lowest single contribution). **m)** Relative abundances of the genus *Akkermansia* in fecal microbial communities before EAE induction. Statistics: One-way ANOVA followed by Tukey’s post-hoc test for groupwise comparison. Non-significant comparisons not shown. *, p < 0.05; **, p < 0.01; ****, p < 0.0001.

Given the limitations of correlation-focused human cohort studies to uncover reliable microbial risk factors for MS susceptibility, the experimental autoimmune encephalomyelitis (EAE) mouse model is commonly used to verify presumed causality between presence of suspected microbial risk factors and development of autoimmune neuroinflammation (*11–13*). However, it is unclear whether the causality of a singular species alone or within only one specific background microbiota, i.e. in mice harboring a relatively consistent, specific pathogen-free (SPF) microbiota composition, can be translateable to the plethora of individual microbiota compositions found across a given population (*1*). Although certain inter-microbial interactions, that promote EAE development, have previously been revealed (*11, 14*), the mutual impact between the background microbiota and potential commensal risk factors on disease-promoting properties of the microbiota is poorly understood.

Here, we investigated whether the EAE disease course can be predicted before disease onset by microbiota-associated readouts. For reliable EAE disease prediction, we weighed in microbial taxonomic composition analyses against microbiota-associated, functional analyses, and we investigated the host immune responses. Our comprehensive approach allowed us to examine how individual host−microbe interactions interfere with disease predictability.

## RESULTS

### *Muc2*-deficiency in mice is associated with less severe experimental autoimmune encephalomyelitis

During experimental autoimmune encephalomyelitis (EAE), the microbiota composition impacts how the host immune system is shaped (*15*), impacting the degree of neuroinfammation. Investigating how different disease phenotypes could be predicted based on the microbiota composition could be a key toward microbiota-based intervention (**Fig. 1a**). As a first step toward testing our hypothesis, we set out to identify the EAE development-associated commensal genera within a complex microbiota. To do so, we induced EAE in specific pathogen free (SPF) mice of different origins and genotypes. We fed these mice diets with different fiber contents, given the impact of dietary fiber quality and quantity on relative abundances of indigenous commensals (*16, 17*). First, we addressed the question of whether changing relative abundances of taxa within a given microbiota might affect outcome of EAE. Toward this goal, we used wildtype C57BL/6J mice purchased from Charles River (CR), which were fed either a standard laboratory chow (fiber-rich; FR) or a fiber-free diet (FF) diet for 20 days, followed by induction of EAE (**Fig. 1b**). Feeding these diets did not result in different disease outcomes (**Fig. 1c–e**), indicating that dietary fiber quantity and quality, and the associated effects on relative abundances of taxa within this particular indigenous microbiota, are not determining factors in mediating EAE (**Fig. 1f**).

Next, we sought to elucidate whether we could observe distinct EAE outcomes between mice whose native microbiota differentiated considerably by the presence of certain taxa, rather than by relative abundances of a shared core microbiota. To do this, we induced EAE in mice deficient for the Muc2 protein (*Muc2*^−/−^), as this genetic modification results in an impaired mucus barrier (*18*). We expected a significantly different indigenous microbiota composition due to anticipated reduction in commensals relying on an intact mucus layer as a functional or nutritional niche (*19*). As controls, we used littermate mice homozygous for the presence of *Muc2* gene (*Muc2*^+/+^). While *Muc2*^+/+^ mice were fed both FR or FF diet, *Muc2*^−/−^ mice were only fed a FR diet (**Fig. 1g**). We observed a significant difference in disease progression between the genotypes, with *Muc2*^−/−^ mice being significantly less susceptible to EAE induction compared to *Muc2*^+/+^ mice, regardless of diet (**Fig. 1h–j**). As observed for CR mice (**Fig. 1f**), diet-mediated influences on disease development were negligeble (**Fig. 1k**).

### Higher abundances of *Akkermansia muciniphila* are associated with less severe EAE in mice with a complex microbiota

To evaluate a potential contribution of the microbiota to the observed differences, we performed 16S rRNA gene-based sequencing analyses on DNA isolated from fecal samples taken before EAE induction (**Fig. 1l–m; Extended Data Fig. 1a–f**) and during the EAE course (**Extended Data Fig. 1a–e**). Intriguingly, the overall microbiota β-diversity (**Extended Data Fig. 1a−d**) and α-diversity (**Extended Data Fig. 1e**) were disconnected from the EAE disease course. Since all four groups of mice expressing the Muc2 protein (CR mice and *Muc2*^+/+^ mice) provided a comparable EAE disease course (**Fig. 1c, h**), which was significantly different from the one observed in *Muc2* knockout (KO) mice (**Fig. 1h**), we assessed potential EAE-relevant microbiota differences by comparing *Muc2*^−/−^ mice (KO) with all *Muc2*-expressing mice combined (WT), irrespective of origin or diet. We identified 11 differentially abundant genera that explained more than 70% of the variance detected in the Bray−Curtis distance matrix between WT and KO mice before induction of EAE (pre-EAE). Pre-EAE relative abundance of the genus *Akkermansia* alone explained 14.4% of said variance (**Fig. 1l**, right panel), correlated negatively with various EAE readouts upon induction of disease (**Fig. 1l** left panel), and was significantly higher in *Muc2*^−/−^ mice compared to WT counterparts (**Fig. 1m**), suggesting possible disease-preventing properties of *Akkermansia*.

However, our observations so far are inadequate to attribute distinct EAE phenoytpes exclusively to changes in *Akkermansia* abundance, or micobiota changes in general, because potential *Muc2* knockout-associated changes in host responses were not specifically addressed. Nonetheless, given that *Akkermansia* is consistently reported as a potential risk factor for MS (*2, 4, 5, 7–9*) and due to its controversial role in EAE development (*2, 20*), we asked whether the observed potential disease-preventing properties in our experiments might be rooted in distinct background microbiota compositions, as *Akkermansia* was embedded in diverse microbiotas with different taxa present or absent (**Extended Data Fig. 1f**).

### Removal of *Akkermansia muciniphila* from a synthetic microbiota results in less severe EAE

To better understand a potential causal role of *Akkermansia* in EAE development and to evaluate its potential as a disease-risk predictor, we colonized germ-free (GF) C57BL/6 mice with a functionally-characterized 14-member human synthetic microbiota (SM14) (*16, 21*) (**Fig. 2a**). This approach allowed us to drop out specific species-of-interest from this community to investigate the contribution of a single microbe on EAE development in a genetically homogenous host. *Akkermansia muciniphila*, the type species for the *Akkermansia* genus, is a member of this SM14 community. Thus, we colonized GF C57BL/6 mice with either the complete SM14 community or a SM13 community, lacking *A. muciniphila*, followed by induction of EAE (**Fig. 2b**). SM13-colonized mice exhibited a significantly less severe EAE phenotype compared to SM14-colonized counterparts (**Fig. 2c**, left panel; **Fig. 2d−f**), highlighting the general contribution of the microbiota to EAE development and the disease-driving role of *A. muciniphila* in the SM14 microbiota-based mouse model when this species is combined with the 13 strains listed in **Fig. 2a**. As controls, we induced EAE in *A. muciniphila*-monoassociated (SM01) and GF mice (**Extended Data Fig. 2**). SM01-colonized and GF mice provided a low-to-intermediate EAE disease phenotype (**Extended Data Fig S2b−d**).

**Figure 2:**
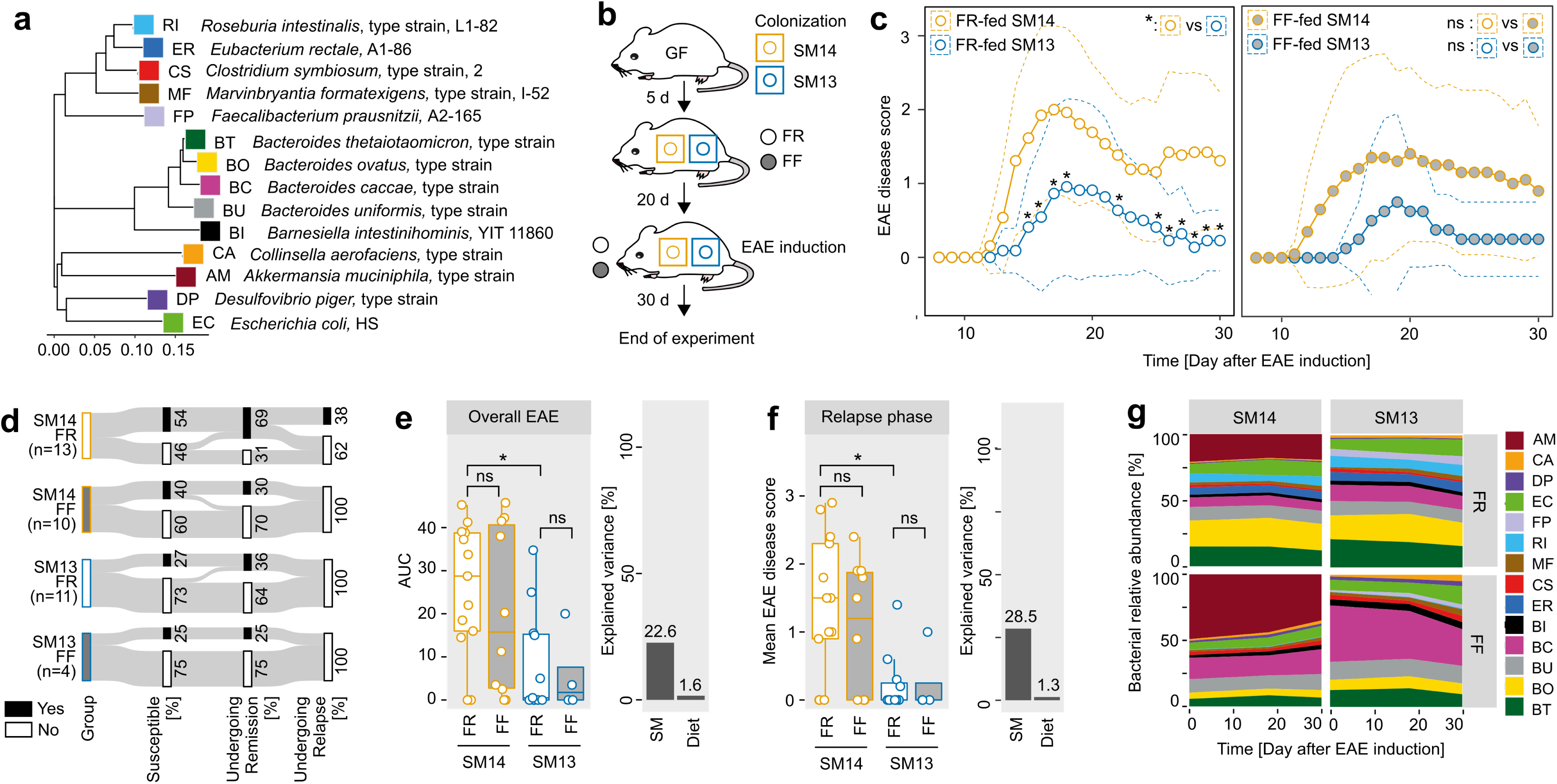
Removal of *Akkermansia muciniphila* from a synthetic human microbiota results in reduced neuroinflammation in EAE-induced mice. **a)** Phylogenetic tree comprising the SM14 community (see Materials and Methods for strain designations and construction of the phylogenetic tree). **b)** Experimental setup. GF C57BL/6N mice were colonized with either SM14 or SM13 (SM14 w/o *A. muciniphila*) communities. 5 d after initial colonization, mice harboring both SM combinations were either switched to a fiber-free (FF) diet or were maintained on a standard chow (FR). 20 d after diet switch, EAE was induced in all mice and disease course was observed for 30 d. **c)** EAE disease scores as a function of time. All mice (FR-fed SM14, n=13 (three independent experients); FR-fed SM13, n=11 (three independent experients); FF-fed SM14, n=10 (two independent experients); FF-fed SM13, n=4) were scored daily at the same time. Left panel: FR-fed mice. Right panel: FF-fed mice. Dotted lines represent SD. Statistics: daily EAE scores were compared using a Wilcoxon rank-sum test. Left panel: * indicates p < 0.05 when comparing SM14-with SM13-colonized mice (FR-fed only). Right panel: “ns” indicates p > 0.05 on any given day when comparing FR-(left panel) with FF-fed (right panel) mice harboring the same SM combinations. **d)** Sankey diagram of key event occurrence (in % of all mice within one group) during EAE. Same event definition as in Fig. 1e. **e)** Left panel: Area-under-the-curve (AUC) analysis of the disease course depicted in c). Each mouse depicted by a separate point. Statistics: One-Way ANOVA followed by Tukey’s post-hoc test. ns, p > 0.05; * < 0.05. Right panel: Percentage of variance explained (η^2^) by diet or colonization (SM combinations, “SM”) when comparing AUC between all 4 groups depicted in the left panel. **f)** Left panel: Mean EAE score during relapse phase (day 26 to day 30). Each mouse depicted by a separate point. Statistics: One-Way ANOVA followed by Tukey’s post-hoc test. ns, p > 0.05; * < 0.05. Right panel: eta-squared analysis as decribed in e). **g)** Streamplots of bacterial relative abundances as a function of time (days after EAE induction). Mean relative abundances per strain, group and timepoint are shown. 2-letter abbreviations of SM14- and SM13-constituent strains are explained in panel a). Bacterial relative abundances were determined by 16S rRNA gene based Illumina sequencing.

To evaluate whether changes in relative abundances of SM14-constituent strains might affect EAE disease course, we fed SM14- and SM13-colonized mice an FF diet, followed by EAE induction (**Fig. 2b**). GF mice were also fed a FF diet to exclude microbiota-independent but diet-mediated effects on EAE. Feeding SM14-colonized mice the FF diet resulted in significantly increased *A. muciniphila* relative abundances compared to equally colonized FR-fed mice (**Fig. 2g**). However, we did not detect any significant differences in any EAE-associated readout (**Fig. 2c**, right panel; **Fig. 2f**) between FR- and FF-fed mice harboring the same microbiota. Removal of *A. muciniphila* from the SM14 community explained between 22% and 28% of the variance for different EAE-associated readouts (**Fig. 2e, f; Extended Data Fig. 2a**, right panel). Given that SM01-colonized mice only provided an intermediate disease phenotype (**Extended Data Fig. 2b−d**), our results suggest that the presence of *A. muciniphila* represents a potential microbial risk factor for severe EAE when combined with certain other strains (SM14) and that changes in its relative abundance within *A. muciniphila*-containing communities negligibly impact EAE disease course.

### *Akkermansia muciniphila*-associated cecal concentrations of γ-amino butyric acid are linked to EAE severity

To evaluate how *A. muciniphila* might alter microbiota function (**Fig. 1a**) within the SM14 reference microbiota, we performed metabolomic and metatranscriptomic analyses. EAE is associated with changes in either plasma metabolite profiles (*22, 23*) or changes in metabolic pathways of the intestinal microbiota (*24*). Furthermore, *A. muciniphila* mediates other pathologies, at least in part, via secretion of certain metabolites (*25*). Thus, we asked whether different levels of neuroinflammation between SM14- and SM13-colonized mice could be explained by *A. muciniphila*-associated metabolite patterns in the cecum or serum. In addition to collecting cecal and serum samples from EAE-induced GF, SM01-, SM13- and SM14-colonized mice, we also collected the same samples from the same groups of non-EAE induced mice. The cecal metabolite profiles were similar between EAE-induced and non-EAE induced groups harboring the same microbiota, as well as between EAE-induced SM13-colonized and SM14-colonized mice (**Fig. 3a, b**). As broader metabolic profiles were disconnected from the EAE disease course (**Extended Data Fig. 3a**), we reasoned that only a few cecal metabolites, if any, might causally influence the EAE disease course.

**Figure 3:**
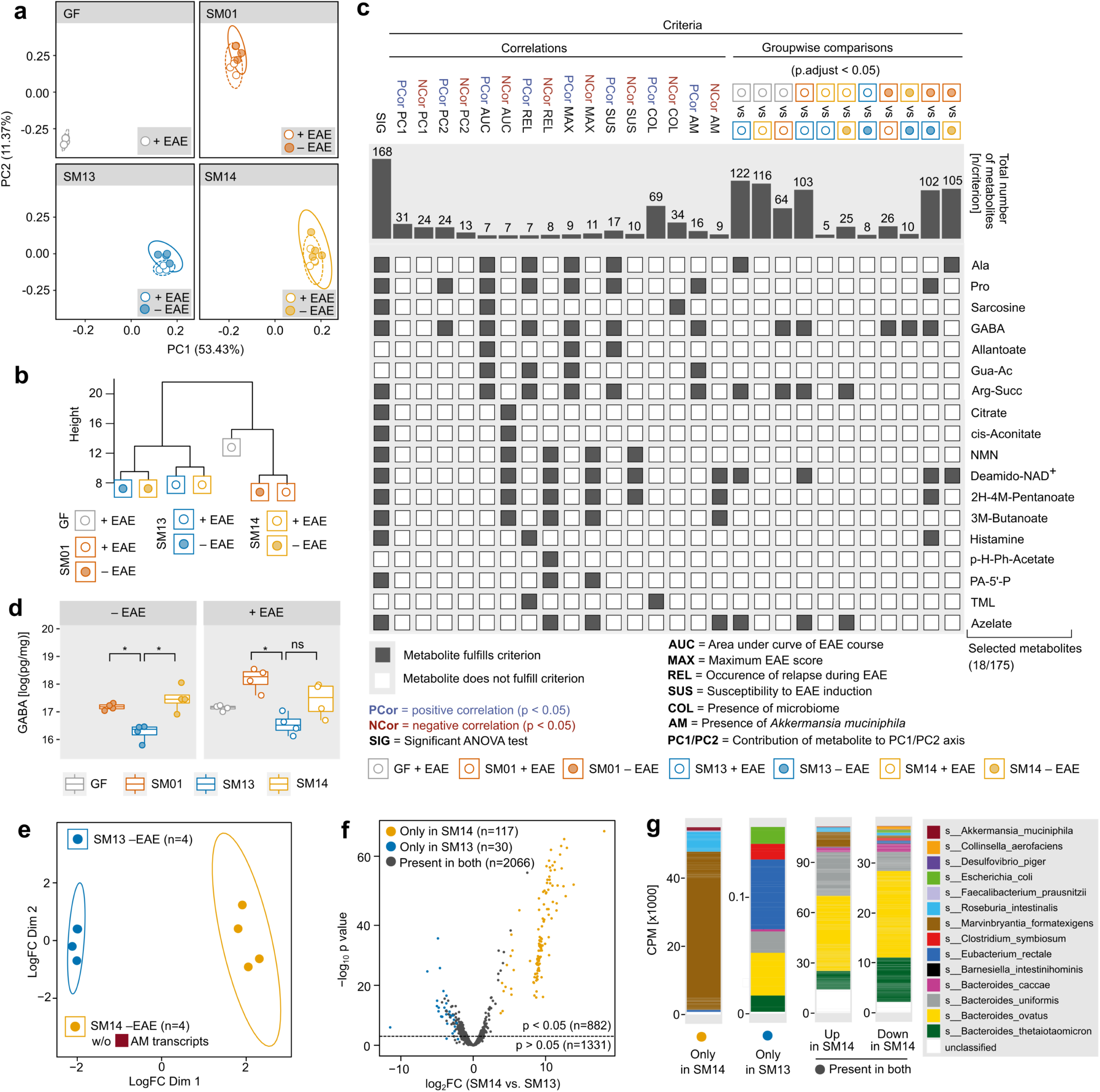
*Akkermansia muciniphila*-mediated autoimmune neuroinflammation is associated with increased cecal concentrations of γ-amino butyric acid. **a–d)** Identification of microbiota-induced and EAE disease course-influencing metabolites using a metabolite screening pipeline. Germ-free (GF) C57BL/6 mice were either colonized with *Akkermansia muciniphila* only (SM01), the 13SM community (SM13), the 14SM community (SM14) or remained GF. 25d after initial colonization, mice were either subjected to harvesting cecal contents (“− EAE”, n = 4 per group) or were EAE-induced followed by cecal content harvest 30d after EAE induction (“+ EAE”, n = 4 per group). Cecal contents were subjected to CE-TOF/MS-based metabolomics analysis. A total of 175 metabolites were identified in at least 50% of the samples in at least one group and were thus included in the overall analysis pipeline. **a)** Principal component analysis (PCA) of log-normalized metabolite concentrations using a Euclidian distance matrix. Panels separated by microbiota compositions. **b)** Hierachical clustering of seven groups based on scaled group means of log-normalized concentrations of each detected metabolite. **c)** “Criteria intersection analysis” of the 175 detected metabolites. Criteria were categorized into “correlation criteria”, summarizing the results of either statistically significant positive (“PCor”) or statistically significant negative Spearman correlation (“NCor”) across all samples of all groups (both, EAE-induced and non-induced mice) and “groupwise comparison criteria”. Correlations referring to EAE-associated criteria (“AUC”: Area under the curve of the disease course; “MAX”: maximum EAE score; “SUS”: susceptibility to EAE induction as defined in Fig 1c; “REL”: occurrence of relapse during the last 5 days of EAE course as defined in Fig 1c) were calculated using samples of EAE-induced mice only. Groupwise comparisons include metabolites found to be significantly different between two given groups based on an unpaired t-tests of log-normalized concentrations, using a Benjamini-Hochberg (BH)-adjusted p-value (p.adjust) as significance criterion. Barplots indicate the total number of metabolites which fulfill each listed criterion. Of the total 175 metabolites, only the criteria intersections of the 18 metabolites that demonstrate a significant correlation with either AUC or REL (or both) are displayed in detail. Grey squares indicate that a given metabolite (on the y-axis) fulfilled a specific criterion (on the x-axis) while white squares indicate a failure to fulfill a given criterion. **d)** Boxplots of log-normalized concentrations of γ-amino butyric acid (GABA) under EAE or non-EAE induced conditions. Statistics were calculated using am unpaired t-test with BH correction for multiple comparisons. **e–g)** Metatranscriptomic analysis of cecal contents of non-EAE-induced SM14-colonized (n=4) and SM13-colonized (n=4) mice. Analysis includes 2213 product-annotated transcripts, surpassing a threshold of 50 CPM in at least 2 samples, accounting for 80–85% of the total CPM. **e)** Multidimensional reduction of transcriptome profiles. In case of SM14-colonized mice, all product-annotated transcripts predicted to be derived from *A. muciniphila* were removed and CPM in these samples were recalculated. This allowed to compare transcriptome profiles of SM13-constituent strains in absence and presence of *A. muciniphila*. **f)** Volcano plot with each dot representing one transcript. Log_2_-transformed ratio of fold-change (FC) relative transcript abundance (Log_2_FC) in SM14-colonized mice compared to SM13-colonized mice shown on the x-axis. p-value of the respective log-transformed FC shown on the y-axis. Dotted line represents significance threshold (p > 0.05). Yellow dots represent product-annotated transcripts only found in SM14-colonized mice (Group CPM mean > 10). Blue dots represent product-annotated transcripts only found in SM13-colonized mice. Grey dots represent product-annotated transcripts found in both groups. n represents number of product-annotated transcripts. **g)** Left 2 panels: Number (n) of genes and cumulative CPM (× 1000) of transcripts being expressed only in either SM13-colonized or SM14-colonized mice. Right 2 panels: Number (n) of genes and cumulative CPM (× 1000) of transcripts being either upregulated or downregulated in SM14-colonized mice, but present in both, SM14- and SM13-colonized mice.

To identify such potential EAE-impacting metabolites, we developed a metabolite-of-interest screening pipeline including 20 independent analyses (**Extended Data Fig. 3b−e**). We proposed that a potential *A. muciniphila*-associated and disease-mediating metabolite should fulfill five different criteria. The rationale for these criteria and the analytical approach is specified in the Materials and Methods section. Among the 18 metabolites that significantly correlated with at least one EAE-associated readout, only γ-amino butyric acid (GABA) emerged as a metabolite-of-interest in cecal samples (**Fig. 3c**). Of note, its concentration was significantly elevated in non-EAE-induced mice harboring an SM combination which resulted in severe EAE upon disease induction. (**Fig. 3d**). Given that GABA concentrations were higher in disease-prone, *A. muciniphila*-harboring mice, these results suggested that the cecal concentration of GABA before EAE induction already defined disease development upon disease induction and that its concentration was linked to disease-influencing properties of the tested microbial communities harboring *A. muciniphila*. Additionally, we did not identify any metabolite-of-interest in serum samples (analyses not shown), indicating that potential metabolite-driven impacts on EAE disease course occur locally in the intestine.

### Presence of *Akkermansia muciniphila* significantly alters gene expression profiles of other microbial community members

Given these community-dependent alterations in microbial metabolite profiles, we rationalized that the presence or absence of *A. muciniphila* had a significant impact on gene expression profiles of the overall microbiota, possibly contributing to distinct EAE phenotypes. Thus, we performed metatranscriptomic analysis of cecal contents obtained from non EAE-induced SM14- and SM13-colonized mice (**Fig. 3e−g**). When comparing transcript profiles of both groups, we found 117 genes expressed only in SM14-colonized mice (**Fig. 3f**). Although we expected that these transcripts would be mostly from *A. muciniphila*, in fact, most of these genes were exclusively expressed by either *Roseburia intestinalis* or *Marvinbryantia formatexigens* (**Fig. 3g**). Of the 30 genes expressed only in SM13-colonized mice, the majority were expressed by *Eubacterium rectale* (**Fig. 3g**). These findings highlight the crucial impact of the presence of a single commensal on the gene expression pattern of other microbial community members, likely impacting their “function” (**Fig. 1a**) within a given community. These indirect influences on community function might also contribute to microbiota-mediated effects on EAE development and thus impact disease-mediating properties of potential risk-predicting species.

### Mucin-degrading capacity of the microbiota is not linked to EAE severity

So far, our results suggest that predicting EAE development, based on the presence or absence of *A. muciniphila*, a mucin-specialist bacterium (*16*), was only possible in mice harboring a variation of the SM14 community (**Figs. 2, 3**) and not in mice harboring a complex community (**Fig. 1**). To further address potential reasons for these discrepancies in general, and the apparent crucial impact of the presence of *A. muciniphila* in SM14-colonized mice in particular, we next hypothesized that changes in relative abundances of other strains in response to dropping out *A. muciniphila* from the SM14 community might causally impact EAE development. We observed that four strains were significantly higher in abundance in SM13-colonized mice compared to SM14-colonized mice (**Fig. 4a**). To address their potential contribution to EAE development, we colonized mice with three additional SM combinations (**Fig. 4b, Extended Data Fig. 4a**). In the first of these combinations, we colonized GF mice with an SM12 community (**Fig. 4b**), lacking *A. muciniphila* and *Faecalibacterium prausnitzii*. This experiment was performed to elucidate whether the >1000-fold increase in relative abundance of *F. prausnitzii* (**Fig. 4a**, extreme right panel) — a species known for gut health-promoting properties (*26*) and decreased abundances in MS patients (*10*) — in mice lacking *A. muciniphila* (**Fig. 4a**) was responsible for EAE-preventing properties of the SM13 community. Intriguingly, SM12-colonized mice (**Fig. 4c−e**) provided a comparable disease course as SM13-colonized mice (**Fig. 2**), suggesting that *F. prausnitzii* expansion in SM13-colonized mice is not responsible for decreased EAE in SM13-colonized mice. At the same time, these data point out the *A. muciniphila*-mediated inhibitory effects on the expansion of an anti-inflammatory bacterium, *F. prausnitzii*.

**Figure 4:**
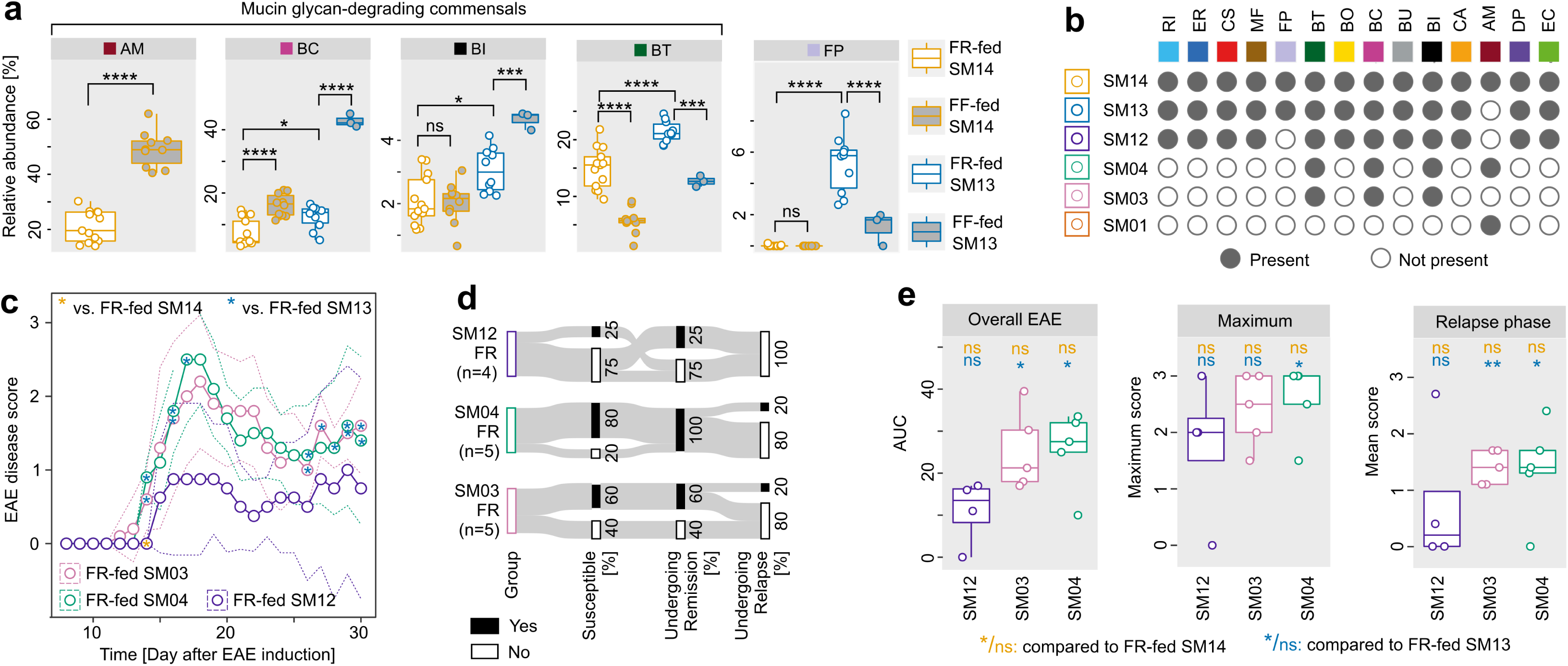
Mucin-degrading capacity of the microbiota is disconnected from EAE disease course. **a)** Data from 16S rRNA gene based Illumina sequencing. Relative abundances of strains, which provided statistically significant differences between FR-fed SM14-colonized mice and FR-fed SM13-colonized mice, as determined by one-way ANOVA followed by Tukey’s post-hoc test, on the day of EAE induction (“before EAE”). *, 0.01 < p < 0.05; **, 0.001 < p < 0.01; ***, p < 0.001. **b)** Constituent strains of all used SM communities. Two-letter abbreviations explained in Fig. 2a. **c)** EAE disease scores as a function of time. All mice (FR-fed SM03, n=5; FR-fed SM04, n=5; FR-fed SM12, n=4) were scored daily at the same time. Statistics: daily EAE scores were compared using a Wilcoxon rank-sum test with BH correction for multiple comparisons. Blue asterisks represent comparison with FR-fed SM13-colonized mice, yellow asterisks with FR-fed SM14-colonized mice. * indicates p < 0.05. **d)** Sankey diagram of key event occurrence (in % of all mice within one group) during EAE. Same event definition as in Fig. 1e. **e)** Left panel: Area-under-the-curve (AUC) analysis of the disease course depicted in c). Each mouse depicted by a separate point. Middle panel: Maximum EAE score per mouse. Right panel: Mean EAE score during relapse phase (d 26 to d 30). Statistics: One-way ANOVA followed by Tukey’s post-hoc test. Blue font indicates comparison with FR-fed SM13-colonized mice, yellow font with FR-fed SM14-colonized mice. *, 0.01 < p < 0.05; **, p < 0.01.

Removal of *A. muciniphila* from the SM14 community further resulted in expansion of three mucin glycan-degrading (*16*) Bacteroidetes species (**Fig. 4a**) in the SM13 community. Thus, we investigated whether colonization with the three mucin glycan-degrading strains alone (SM03) resulted in decreased EAE compared to SM14-colonized mice and whether addition of *A. muciniphila* (SM04) might counteract a potential beneficial effect. While SM03- and SM04-colonized mice showed comparable EAE disease courses (**Fig. 4c−e**) to SM14-colonized mice (**Fig. 2**), they differed significantly from SM13-colonized mice. Additionally, the three mucin glycan-degrading Bacteroidetes strains appeared to not provide disease-reducing properties but, on the contrary, disease-promoting properties in the absence of the remaining 10 strains within the SM13 community. To evaluate whether dysregulated mucin turnover might contribute to the observed results in these mice, we assessed various indirect measures for intestinal barrier integrity. We did not detect any correlations between EAE outcome and glycan-degrading enzymatic activities (**Extended Data Fig. 4b−g**), serum concentrations of LPS, occludin or zonulin as well as with fecal concentrations of lipocalin (**Extended Data Fig. 4h,i**), or SCFA (**Extended Data Fig. 4j**). Thus, our results suggest that bacterially-mediated mucus glycan degradation or barrier integrity impairment, in the context of the microbiota combinations used in this study, were not an individual predictor for EAE disease development.

### Microbiota composition can be used to estimate the probability of severe EAE incidence

Thus far, groupwise comparisons of EAE-associated readouts and microbiota compositions failed to identify reliable predictors for disease development in EAE-induced mice. Therefore, we next aimed to elucidate common denominators on a group-based and individual level to help uncover more reliable potential predictors for microbiota-mediated impacts on disease course. First, we conducted group-based comparison of EAE outcomes between all 10 tested diet−colonization combinations (“groups”) (**Fig. 5a, b**). Performing hierarchical clustering (**Fig. 5c**) based on group means of key EAE-associated readouts (**Fig. 5b**) revealed three distinct group phenotypes: “moderate”, “intermediate” and “severe”. While diet explained less than 8% of the variance observed for EAE-associated readouts, microbiota composition (SM) explained between 11% and 27% (**Fig. 5d**). Given these low values, rooted in considerable intra-group variances (**Fig. 5b**), we performed individual EAE phenotype clustering, treating all mice across all groups individually (**Fig. 5e**). T-distributed stochastic neighbor embedding (t-SNE) analysis of all EAE-induced individuals resulted in two disease clusters: “Cluster 1”, comprising mice showing strong EAE symptoms, and “Cluster 2”, comprising mice showing minor EAE symptoms (**Fig. 5e**). Besides SM03- and SM04-colonized mice, every group included mice of both phenotypes (**Fig. 5f**), however with varying proportions. These proportions broadly, but not completely, corresponded to the group-based phenotype classification (**Fig. 5c**). In summary, these results (**Fig. 5a−f**) indicate that knowing the composition of the microbiota, in combination with information on dietary conditions, enables estimation of the probability for either moderate or severe disease, but is unsuitable to predict individual EAE outcomes.

**Figure 5:**
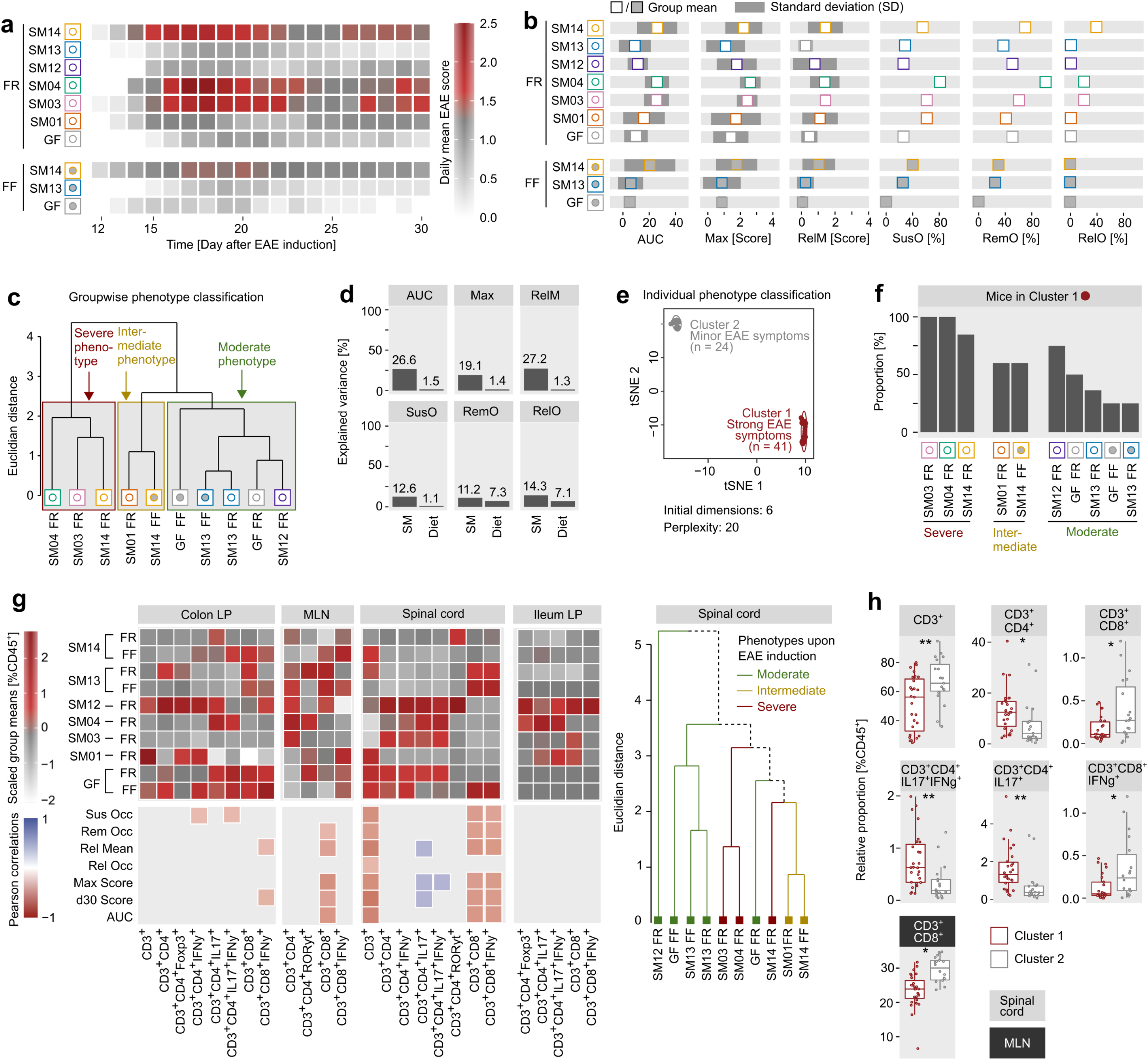
Microbiota alterations result in three different EAE group phenotypes and two different individual phenotypes. **a)** Summary of EAE disease course of all tested colonization/diet-combinations. Heatmap summarizes data from Fig. 2c, Extended Data Fig. 2 and Fig. 4c. Daily mean EAE score per colonization/diet-combination visualized by color scale. **b)** Summary of key EAE-associated readouts of all tested colonization/diet-combinations. Squares indicate group mean. Dark grey bar indicates standard deviation (SD). SusO, occurrence of susceptibility; RemO, occurrence of remission; RelM, mean EAE score during relapse phase; RelO, occurrence of disease relapse; Max, maximum achieved EAE score; AUC, area-under-the-curve analysis of EAE score as a function of time. For definition of “Susceptibility”, “Relapse” and “Remission” see Fig. 1e. **c)** Groupwise EAE phenotype classification. Cluster dendrogram of scaled group means of EAE-associated readouts (panel b), based on a Euclidian distance matrix. Group phenotypes were classified according to the three obtained main clusters. **d)** Percentage of variance explained (η^2^) by diet or SM-combinations (SM) when comparing EAE-associated readouts among all colonization/diet-combinations. **e)** EAE phenotype classification by mouse (individual) by applying t-stochastic neighbor embedding (t-SNE) analysis to EAE-associated readouts data sets of each individual mouse across all tested colonization/diet-combinations. Cluster 1 (strong EAE symptoms) and Cluster 2 (minor EAE symptoms) phenotype) resulted from applying a perplexity of 20 to t-SNE analysis, using 6 initial dimensions. **f)** Proportion of mice in Cluster 1 per colonization/diet-combination. Groupwise EAE phenotype classification (panel c) added below. **g)** Upper panel: Relative abundances of T cell subsets [%CD45^+^], scaled by subset and organ, of those subsets which provided significant differences as determined by one-way ANOVA. Lower panel: Correlations between relative abundances of those subsets with key EAE-associated readouts in the same individuals. All mice of all groups included irrespective of microbiota composition. Right: Hierarchical group clustering of all 10 tested colonization-diet combinations based on subset means of significantly different T cell subsets in the spinal cords. **h)** Relative abundances of T cell subsets, which were found to be statistically significant by EAE phenotype cluster affiliation, as determined by unpaired t-tests. *, 0.01 < p < 0.05; **, p < 0.01.

Given that IL-17- and IFNγ-producing CD4^+^ cells (*11*), CD8^+^ cells (*27*), and IgA^+^ IL-10^+^ plasma cells (*28*) are reported to link the microbiota with EAE development, host-specific influences on these immune responses might contribute to individually different disease outcomes despite identical microbiome compositions. Examining such effects in more detail is critical for making disease course predictions based on microbiota composition or function. Thus, we analyzed T and B cell polarization in mice before and after EAE induction to elucidate whether potential host-specific responses occurred upstream or downstream of immune cell activation. While we detected no differences in B cell subsets among EAE-induced mice, we found 25 T cell populations in four different organs with significantly different relative abundances between groups (**Fig. 5g**, left panel; **Extended Data Fig. 5a**). Among those, nine populations also correlated with individual outcome of EAE (**Fig. 5g**, left panel). Among all tested organs (colonic and ileal lamina propria, mesenteric lymph nodes and spinal cords), relative abundances of T cell populations in the spinal cords corresponded best, but not perfectly, to the respective EAE group phenotype (**Fig. 5g**, right panel). While group-wise comparison of each T cell population failed to explain observed EAE phenotypes, individual cluster-based analyses (**Extended Data Fig. 5e**) showed significant differences for seven populations, with IFNγ-expressing Th17 cells in the spinal cords significantly increased in Cluster 1-mice (**Fig. 5h**). Mouse-specific T cell polarization profiles aligned better with disease outcome (**Fig. 5h**) than with SM−diet combinations (**Fig. 5h**). Thus, we concluded that host-specific differences must occur before T cell activation in EAE-induced mice, most probably due to individual microbiota-mediated signals that appeared to be distinct even in mice harboring the same set of strains.

When analyzing T cell subsets in non-EAE induced mice, we found that the microbiota composition primed CD4^+^ T cells towards a pro-inflammatory Th17 response before EAE-induction (**Extended Data Fig. 5b**). Although we found more significant correlations of these populations with EAE-associated readouts in the ileum, overall T cell population distribution in the colon aligned best with emerging EAE group phenotypes upon EAE induction (**Extended Data Fig. 5c,d**), suggesting a crucial contribution of T cell priming in the colon by the microbiota to disease development upon EAE induction.

In summary, it was impossible to predict individual disease development based on microbiota−diet combinations alone despite apparent microbiota-mediated priming of the local adaptive immune system before EAE induction. This observation suggests that microbiota-mediated signals that influence the adaptive immune system to either promote or decelerate EAE development are relatively constant before disease induction, but are prone to individual changes upon disease onset.

### IgA coating index of *Bacteroides ovatu*s represents a surrogate measure to predict microbiota-mediated impact on individual EAE severity

Considering these individual signaling changes, we next targeted identification of microbiota-associated factors suitable to predict individual outcome of EAE. For each given strain (**Fig. 6a**), we first assessed whether its relative abundance before EAE induction (**Extended Data Fig. 6a**), as determined by 16S rRNA gene sequencing in each individual mouse, allowed for prediction of individual disease course after EAE induction (**Fig. 6b, c**). To do so, we only included mice harboring at least 12 different strains, thus excluding mice with low-diversity microbiotas (SM04, SM03, SM01). Correlations for each strain were only assessed for those mice, which were gavaged with the respective strain and calculations were performed by either including mice harboring SM14, SM13 and SM12 communities, or specific combinations thereof, into the analysis. We found statistically significant correlations between pre-EAE bacterial relative abundances with EAE-associated readouts for some strains (**Fig. 6b)**. However, the few statistically significant correlations we determined were generally weak (R<0.4) and Pearson correlation values for a given strain were highly dependent on the background microbiota (**Fig. 6b)**. In line with this, relative abundances of strains only explained very low proportions of the variances across all groups for all assessed EAE-associated readouts (**Fig. 6c**).

**Figure 6:**
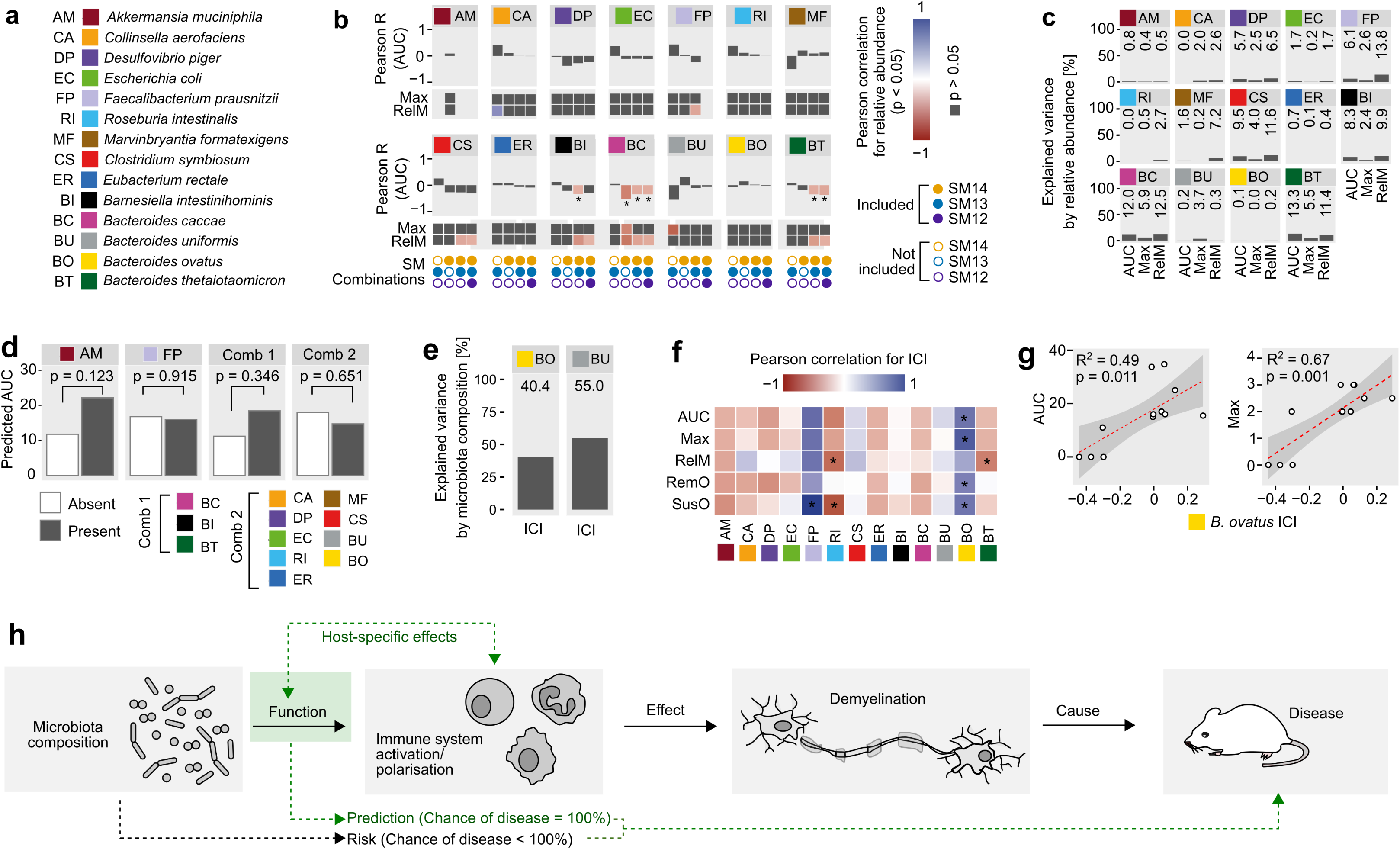
Using IgA coating index of an inert reporter species as surrogate measure to assess EAE-mediating properties of a given microbial community. **a)** Color codes and 2-letter abbreviations of all SM14-constituent strains **b)** Pearson correlation between bacterial relative abundance of each strain before EAE induction, as detected by 16S rRNA gene sequencing, with several EAE-associated readouts for individual mice: AUC, area-under-the-curve analysis of EAE score as a function of time; Max, maximum achieved EAE score; RelM, mean EAE score during relapse phase. For AM, only SM14-colonized mice could be used for correlation analysis. For FP only SM13- and SM14-colonized mice could be used. For all other strains, SM14-, SM13-, and SM12-colonized mice were analysed. Mice of both dietary groups (FR and FF) were included in this analysis. Pearson correlation values for association between bacterial relative abundance and AUC are shown as bar plots, for Max and RelM only color-coded squares are shown. Significant correlations (p < 0.05) are shown in colors from blue to red according to the legend; insignificant correlations are shown in dark grey only. Correlations for each strain were calculated for four different microbiota (SM) combinations: SM13 only; SM14 only; SM13 and SM14 combined; SM12, SM13 and SM14 combined. **c)** Variance of three key EAE-associated readouts explained by relative abundance of a given strain before EAE induction. Analysis performed by combining SM12-, SM13- and SM14-colonized mice irrespective of fed diet. **d)** Linear mixed model regression for predicted AUC with presence of the strain as an independent variable and colonization as a random intercept effect. **e)** Variance in IgA-coating index (ICI) explained by high-diversity background microbiota compositions (SM12, SM13 and SM14) in two strains providing highest microbiota-dependencies on ICI. **f)** Individual-based paired Pearson correlation of key EAE associated readouts with ICIs of SM14-constituent strains before EAE induction. Mice of all colonization-diet combinations were included in this analysis. *, p < 0.05 (without further distinction). **g)** Correlation of *B. ovatus* ICI with AUC (left) and maximum EAE score (right) in all mice harboring *B. ovatus*, irrespective of background microbiota. **h)** Graphical summary: Taxonomic microbiota information can be used to assess the “Risk” of a given individual to develop disease (as defined by a chance < 100 %). For “Predictions”, as defined by a 100 % chance to develop disease, host-mediated influences on the microbiota function within a given individual must have to be taken into account.

Next, we asked whether presence or absence of a given strain might be a better predictor for individual EAE development. Thus, we performed a linear mixed model regression for three EAE-associated readouts with presence of the strain as an independent variable and colonization as a random intercept effect (**Fig. 6d**, **Extended Data Fig. 6b**). Given the setup of our tested SM-combinations, we could only assess *A. muciniphila* and *F. prausnitzii* separately and had to analyze the remaining 12 strains in groups of two combinations. Presence or absence of a specific strain or strain combination was insufficient to predict the individual outcome of any of the tested EAE-associated readouts (**Fig. 6d, Extended Data Fig. 6b**), indicating that potential disease-driving or -preventing properties of a given strain or strain combination is crucially determined by the background microbiota.

Coating of intestinal commensals by host plasma cell-derived IgA represents a crucial host response for maintaining immune homeostasis in the context of autoimmune neuroinflammation (*29–31*). Secretory IgA (sIgA) levels in the mouse feces were disconnected from individual EAE outcomes (**Extended Data Fig. 6c**), but were strongly connected to the microbiota composition (**Extended Data Fig. 6d**). Interestingly, we found a significant correlation between group means of sIgA concentrations and corresponding EAE susceptibility incidence (**Extended Data Fig. 6e**). Owing to these observations and given that the “IgA-coating index” (ICI) was previously suggested to be a measure of autoimmunity-promoting potential of a given commensal species (*32*), we determined ICIs for each strain within each high-diversity SM combination (SM12, SM13 and SM14) in every individual (**Fig. 6e−g, Extended Data Fig. 6f−h**). Interestingly, there was a tendency for ICIs to differ between distinct SM-combinations **(Extended Data Fig. 6f)** and microbiota composition explained more than 40% of the variance between ICIs of *B. ovatus* and *B. uniformis* **(Fig. 6e, Extended Data Fig. 6g)**, suggesting a crucial role of the background microbiota on strain-specific IgA coating (**Extended Data Fig. 6h)**. Additionally, ICIs of these strains varied not only between distinct groups, but also between individuals within groups. Thus, we reasoned that the individual ICI of these strains might reflect individual EAE-promoting properties of the microbiota in a certain host. Correlation analysis of strain-specific ICI, as determined from fecal samples obtained before EAE induction, with EAE outcome in the same individual, revealed significant correlations with some EAE-associated readouts for four strains (**Fig. 6f**). However, the only strain whose individual ICI provided significant correlations with the two most important EAE-associated readouts (AUC and maximum achieved EAE score) was *B. ovatus* (**Fig. 6f, g**), thus allowing for individual prediction of EAE disease course across all *B. ovatus*-encompassing SM combinations (**Fig. 6g**).

## DISCUSSION

Given the association between the intestinal microbiota and extra-intestinal autoimmune diseases (*1*), microbiota manipulation is a feasible approach to boost existing therapy options for MS patients. Potential strategies for microbiota modulation include administration of antibiotics or probiotics (*33*), dietary interventions (*34, 35*) or fecal microbiota transplantation (*36*). However, such untargeted strategies could lead to broad-scale changes in the microbiota with potentially unpredictable outcomes. On the contrary, given the unique microbiota composition in a given individual, personalized approaches might be more promising.

A better understanding of what exactly makes a specific microbiota composition in a given individual disease prone is a precondition for a targeted personalized approach. Such knowledge is expected to result in analytical procedures to evaluate the average risk of disease or even predict individual outcomes. Previously suggested measures to evaluate the MS-mediating risk of a microbial community, such as the microbiota α-diversity (*37*) or the Firmicutes-to-Bacteroidetes ratio (*38*), emerged as unsuitable tools (*10*) and more focus is currently being put on individual taxa or combinations of taxa (*10*). A recent mouse study suggested that up to 50 different microbial taxa could be associated with disease (*39*). Disease-associated bacterial taxa are often referred to as “microbial risk factors”, which are mostly identified by differences in presence or relative abundances from cross-sectional human cohort studies. *Akkermansia muciniphila* represents such a potential microbial risk factor as it was reportedly increased in MS patients across various human cohorts (*2, 5, 7, 40*). Other studies, however, report on positive effects of *A. muciniphila* on maintaining general gut homeostasis (*41, 42*) or on progression of autoimmune neuroinflammation in mice (*20*).

At first sight, these observations might appear contradictory. They are, however, corroborated by our findings. By comparing development of experimental autoimmune encephalomyelitis (EAE) in mice of different genetic backgrounds and with distinct complex microbiotas, we found the genus *Akkermansia* to be the most negatively associated with EAE disease development, thus representing a potential hallmark genus for less severe EAE, when considering the microbiota composition as the only variable and ignoring host genetics (**Extended Data Fig. 7**). Next, we evaluated whether this finding could be reproduced in gnotobiotic, genetically homogenous mice harboring different combinations of a reduced reference microbiota, with or without *A. muciniphila*. Interestingly, we found *A. muciniphila* to be positively associated with EAE severity in certain mice harboring specific reduced communities. This was associated with increased cecal levels of γ-amino butyric acid (GABA). However, we cannot reliably conclude whether increased GABA concentrations are a general risk factor for EAE induction or whether this only applies to a certain microbiota composition. Since elevated GABA levels were previously reported to be associated with less neuroinflammation (*43, 44*), we deemed assessment of intestinal GABA concentration, without corresponding information on microbiota composition, unsuitable for disease course prediction. It is unclear whether this neurotransmitter directly mediates EAE-influencing host responses via interaction with local host receptors (*45*) or whether it might act as a signaling molecule or energy source (*46*) for other species, which finally mediate disease promotion.

Our study suggests that focusing on microbiota composition only, as we did in our experiments using mice of different genotypes, can result in misleading conclusions. Co-variates, such as diet, sex, medication use, geographic location, disease subtypes and genetic heterogeneity of study participants make interpretation of microbiota data from human cohort studies complicated, although certain biostatistic approaches help to reduce the risk of misinterpretation, as elegantly shown in a recent publication from the iMSMS Consortium (*10*). In addition, we found that 16S rRNA gene sequencing-based determination of relative taxa abundances is unsuitable to make meaningful assumptions on disease-mediating properties of a given microbiome. Assessing the presence or absence of taxa allowed us to determine the probability of severe disease (**Fig. 6h**: “Risk of disease”). However, this observation was unrelated to presence or absence of a single taxon, suggesting that focusing on combinations of taxa and/or environmental factors, rather than single taxa alone, may be required to form reliable conclusions. Thus, our results suggest that mutual influences between a suspected risk factor and the microbial environment crucially shape the overall microbiota’s disease-impacting potential **(Extended Data Figure 7)**.

Our metatranscriptomics analyses revealed that even minor changes in microbiota composition, i.e. by removing *A. muciniphila* from a reduced community, resulted in profound changes in gene expression patterns of some, but not all, intestinal microbes. Such influences may also affect disease-mediating properties of the microbiota. Therefore, we suggest to put more focus on microbial network analysis to disentangle specific inter-microbial interactions. Although not yet a technically and analytically refined approach (*47*), metagenomic-based microbiota network analyses are currently being explored as an analysis tool (*48*) and might be superior to statistical analysis of species−species co-abundances (*10*). A key study, evaluating the effects of multiple defined microbiota compositions on fitness of *Drosophila melanogaster*, already pointed out that microbial network interactions are more important than relative abundances of a given species alone (*49*). Our study documents similar innovative findings, based on comprehensive datasets, in a controlled vertebrate gnotobiotic disease model..

In addition to these microbiota-specific effects, host-specific effects appeared to be a decisive factor for individual EAE development in our experiments, further complicating the quest for reliable disease predictors. Even in genetically homogenous mice of the same sex and age, harboring the exact same set of commensal bacteria and living under the same standardized conditions, we found considerable individual differences in EAE disease course, suggesting that the individual disease development is mediated by either microbe−microbe or microbe−host interactions (*50, 51*) (**Fig. 6h**). After extensive evaluation of multiple microbiota-associated readouts, we found the IgA-coating index (ICI) of the fiber-degrader *Bacteroides ovatus* (*16*) to be capable of sensing the individual EAE-influencing properties of the microbiota, irrespective of its definite composition. Determining the ICI of *B. ovatus* before disease induction correctly predicted EAE outcome in every individual (**Fig 6h**: “Prediction of disease”). Thus, we propose that *B. ovatus* acts as a “reporter species”, reflecting the individual microbiota- and host-mediated dual influences on EAE progression while taking into account distinct microbiota functions across different hosts (**Fig. 6h**: “host-mediated effects on Function”). In the current study, although such a property of *B. ovatus* was only evaluated in reduced microbial communities, the concept of “reporter species” might also apply to other strains and more complex communities, including MS patient microbiotas. A recent study showed that different strains of *B. ovatus* are capable of driving variable host IgA secretion (*52*), which might also impact the IgA coating of *B. ovatus*. Although we did not evaluate in our study whether different strains of *B. ovatus* evolve in different mice, future studies need to consider such aspects.

In summary, we demonstrate that making disease-course predictions based on microbiota characteristics is generally possible, but is not as black-and-white as it might appear. We therefore strongly argue for a reconsideration of how microbiota-related data are analyzed and interpreted. In particular, we advocate for higher analytical standards, with more sophisticated data integration to better account for discrepancies in host-specific microbiota function.

## MATERIALS AND METHODS

### Institutional Review Board Statement for conduction of mouse experiments

All mouse experiments followed a two-step animal protocol approval procedure. Protocols were first evaluated and pre-approved by either the ethical committee of the University of Luxembourg (AEEC) or the Animal Welfare System (AWS) of the Luxembourg Institute of Health, followed by final approval by the Luxembourgish Ministry of Agriculture, Viticulture, and Rural Development (Protocol numbers: LUPA2020/02, LUPA2020/27, LUPA2020/32, LUPA2019/43, LUPA2020/22, LUPA2019/51) before start of experiments. All experiments were performed according to the Federation of European Laboratory Animal Science Association (FELASA). The study was conducted according to the “Règlement grand-ducal du 11 Janvier 2013 relatif à la protection des Animaux utilisés à des fins Scientifiques” based on the “Directive 2010/63/EU” of the European Parliament and the European Council from September 2010 on the protection of animals used for scientific purposes. All animals were exposed to 12 hours of light daily.

### Origin of mice and housing conditions

For gnotobiotic experiments, female germ-free (GF) C57BL/6N mice were purchased from Taconic Biosciences, USA, at the age of 4 to 8 weeks. All animals were housed and bred in the GF facility of the University of Luxembourg, supervised by the ethical committee of the University of Luxembourg (AEEC). Mice were randomly allocated to different experimental groups and were housed in ISO-cages in groups with a maximum of five animals per cage. Water and diets were provided ad libitum. Before start of experiments, GF status of all mice was confirmed via anaerobic and aerobic incubation of fecal samples in 5 ml culture tubes containing two different rich media, Brain Heart Infusion Broth and Nutrient Broth.

For experiments performed under specific pathogen-free (SPF) conditions, as shown in Fig. 1, we used female mice of different origin. C57BL/6J wildtype mice were purchased from Charles River at the age of 5 to 8 weeks. Furthermore, we used mice lacking the *Muc2* gene (strain designation: 129P2/OlaHsd×C57BL/6-Muc2), which were originally obtained from University of Bern, Switzerland under GF conditions. GF 129P2/OlaHsd×C57BL/6-Muc2 mice were mated with SPF-housed C57BL/6 mice obtained from Charles River resulting in offspring heterozygous for presence of the *Muc2* gene (*Muc2*^+/−^). *Muc2*^+/−^ mice were constantly kept under the same SPF conditions, as the SPF-housed parental C57BL/6 mice. Next, male and female *Muc2*^+/−^ mice were mated and offsprings were genotyped for absence and presence of the *Muc2* gene. Homozygous *Muc2*^−/−^ and *Muc2*^+/+^ mice obtained from this breeding were then used for experiments.

### Genotyping for presence or absence of the *Muc2* gene

Genotyping for presence or absence of the *Muc2* gene from mouse ear tissue was performed using the SampleIN™ Direct PCR Kit (HighQu, #DPS0105) according to the manufacturer’s instructions. Three different primers were used at a final concentration of 0.4 µM in the PCR reaction (Primer sequences, 5’→3’; Primer 1: TCCACATTATCACCTTGAC; Primer 2: GGATTGGGAAGACAATAG; Primer 3: AGGGAATCGGTAGACATC). The PCR was conducted with an annealing temperature of 56 °C and 37 cycles. Presence of the *Muc2* gene resulted in an amplificate of 280 bp, while absence of the *Muc2* gene resulted in an amplificate of 320 bp. Amplificates were visualized on a 1.5% agarose gel by gel electrophoresis.

### Colonization of germ-free mice with a reduced human synthetic microbiota

All 14 bacterial strains of the synthetic microbiota (SM) were cultured and processed under anaerobic conditions using a Type B vinyl anaerobic chamber from LabGene, Switzerland, as published in detail previously (*21*). A total of six different SM combinations were used to colonize GF mice. Non-colonized GF mice were used as a control group. Intragastric gavage and verification of proper colonization of administered strains was performed as described in detail elsewhere (*21*). Details on the 14 different used strains are summarized in **Extended Data Table 1**.

### Mouse diets

Mice were either maintained on a standard mouse chow (fiber-rich; “FR”) or switched to a fiber-free (“FF”) diet. We used two different FR diets with approximately 15% dietary fiber: SAFE A04 chow (Augy, France, product code U8233G10R) for gnotobiotic mice, sterilized by 25 kGy gamma irradiation, and SDS Standard CRM (P) Rat and Mouse Breeder and Grower diet (Special Diets Service; Essex, UK; product code 801722), sterilized by 9 kGy gamma irradiation, for SPF-housed mice. The FF diet was used for both gnotobiotic and SPF-housed mice and was custom-manufactured by SAFE (Augy, France), based on a modified version of the Harlan TD.08810 diet, as described previously (*53*).

### Experimental timeline of mouse experiments

*Experiments performed under gnotobiotic conditions*: At the age of 4 to 8 weeks, mice were colonized with various SM combination (see above), being fed an FR diet. 5 days after initial colonization, mice were either maintained on an FR diet or switched to an FF diet until the end of experiment. Mice were then either induced with experimental autoimmune encephalomyelitis (labelled as “+EAE” in the manuscript) 20 days after the initial gavage or were euthanized for organ harvest 25 days after the initial gavage without induction of experimental autoimmune encephalomyelitis (labelled as “−EAE” in the manuscript). *Experiments performed under SPF conditions*: Mice of all genotypes were raised and maintained on an FR diet. At the age of 6 weeks, mice were either switched to an FF diet or kept on the FR diet. 20 days later, mice fed either diet were subjected to induction of experimental autoimmune encephalomyelitis (EAE). Course of EAE under both, gnotobiotic or SPF, conditions was observed for 30 days.

### Experimental autoimmune encephalomyelitis (EAE)

Mice were immunized using the Hooke Kit™ MOG_35-55_/CFA Emulsion PTX (Hooke Laboratories, #EK-2110) according to manufacturer‘s instructions. In brief, mice were immunized with a subcutaneous injection of a myelin oligodendrocyte glycoprotein-derived peptide (MOG_35-55_) and complete Freund’s adjuvant (CFA) delivered in pre-filled syringes. Subcutaneous injection of two times 100 µL (200 µL in total) of MOG/CFA (1 mg mL^−1^ MOG_35-55_ and 2-5 mg mL^−1^ killed mycobacterium tuberculosis H37Ra/mL (CFU)) mixture was performed on two sides bilateral in each of the mouse’s flank. Additionally, Pertussis toxin (PTX) solution was injected on the day of MOG peptide immunization and 48 hours after the first injection. Glycerol-buffer stabilized PTX was diluted in sterile PBS for application of 400 ng PTX (gnotobiotic experiments) or 150 ng (experiments performed under SPF conditions) by intraperitoneal injection of 100 µL PTX solution. The EAE clinical symptom scores were assessed daily according to the scheme depicted in **Extended Data Figure 8**. For EAE scoring, names of synthetic microbiota groups (such as SM14, SM13, etc.) were blinded. EAE scoring was performed blinded by one researcher (Alex Steimle) and independently unblinded (for microbiota groups) by another researcher (Mareike Neumann), but at the same time. In case of discrepancy of assigned scores by two researchers, the scoring persons discussed and agreed on a certain score. Note that the discrepancy in scoring the same mouse by 2 persons was uncommon due to the rigorous and straight-forward scoring approach outlined in Extended Data Figure 8. If discrepancy happened, the discrepancy was for a max difference of 0.5 score. During EAE scoring, proper care was taken to avoid potential contamination of the gnotobiotic mice.

### Euthanisia and organ harvest

Mice of all groups (“− EAE” and “+ EAE”) were subjected to terminal anesthesia through intraperitoneal application of a combination of midazolam (5 mg kg^−1^), ketamine (100 mg kg^−1^), and xylazine (10 mg kg^−1^) followed by cardiac perfusion with ice-cold PBS. Colonic content, cecal content, blood, and organs were harvested for downstream analysis. Mesenteric lymph nodes (MLN) were removed and homogenized by mechanical passage through a 70 µm cell strainer. MLN cells were washed once in ice-cold PBS for 10 min at 800 × g, resuspended in ice-cold PBS and stored on ice until further use. Colon, ileum were removed and temporarily stored in modified HBSS (“HBSS (w/o)”; Hank’s balanced salt solution buffered without Ca^2+^ and Mg^2+^, buffered with 10 mM HEPES) on ice, while removed spinal cords were temporarily stored in D-PBS. All three organs were then subjected to lymphocyte extraction as decribed below.

### Lymphocyte extraction from colonic lamina propria, ileal lamina propria and spinal cords

After organ removal, lymphocytes from the colonic lamina propria (CLP), ileal lamina propria (ILP) and spinal cords (SC) were extracted. While CLP and ILP lymphocytes were extracted using the lamina propria dissociation kit (Miltenyi Biotec, #130-097-410), SC lymphocytes were extracted using a brain dissociation kit (Miltenyi Biotec, #130-107-677), according to the manufacturer’s instructions. In brief, colon and ileum were dissected and stored in HBSS (w/o). Feces and fat tissue were removed, organs were opened longitudinally, washed in HBSS (w/o), and cut laterally into 0.5 cm long pieces. Tissue pieces were transferred into 20 mL of a predigestion solution (HBSS (w/o), 5 mM EDTA, 5% fetal bovine serum (FBS), 1 mM dithiothreitol) and kept for 20 min at 37 °C under continuous rotation. Samples were then vortexed for 10 sec and applied on a 100 µm cell strainer. Last two steps were repeated once. Tissue pieces were then transferred into HBSS (w/o) and kept for 20 min at 37 °C under continuous rotation. After 10 seconds of vortexing, tissue pieces were applied on a 100 µm cell strainer. Tissue pieces were then transferred to a GentleMACS C Tube (Miltenyi Biotec, #130-093-237) containing 2.35 mL of a digestion solution and homogenized on a GentleMACS Octo Dissociator (Miltenyi Biotec, #130-096-427, program 37C_m_LPDK_1). Homogenates were resuspended in 5 mL PB Buffer (phosphate-buffered saline (PBS), pH 7.2, with 0.5 % bovine serum albumin), passed through a 70 µm cell strainer and centrifuged at 300 × g for 10 min at 4 °C. Cell pellets were resuspended in ice-cold PB buffer and stored on ice until further use. Spinal cords were stored in ice-cold D-PBS (Dulbecco’s phosphate-buffered saline with calcium) until they were transferred to a GentleMACS C Tube containing a digestion solution. Samples were processed on a GentleMACS Octo Dissociator (program 37C_ABDK_01) and rinsed through a 70 μm cell strainer. The cell suspension flowthrough was then centrifuged at 300 × g for 10 min at 4 °C. Debris removal was performed by resuspending the cell pellet in 1550 µL D-PBS, adding 450 µL of Debris Removal Solution, and overlaying with 2 mL of D-PBS. Samples were centrifuged at 4 °C at 3000 × g for 10 min. The two top phases were aspirated and the cell suspension was diluted with cold D-PBS. Samples were then inverted three times and centrifuged at 4 °C, 1000 × g for 10 min and the cell were resuspended and stored in ice-cold D-PBS until further use.

### Cell stimulation and flow cytometry

10^6^ cells (MLN cell suspensions as well as lymphocyte extracts from CLP, ILP and SC) were resuspended in 1 mL complete cell culture medium (RPMI containing 10% FBS, 2 mM Glutamine, 50 U × mL^−1^ penicillin, 50 µg × mL^−1^ streptomycin, and 0.1% mercaptoethanol) supplemented with 2 μL Cell Activation Cocktail with Brefeldin A (Biolegend, #423304) and incubated for 4 h at 37 °C. Cells were centrifuged at 500 × g for 5 min, resuspended in 100 μL Zombie NIR (1:1000 in PBS, Zombie NIR™ Fixable Viability Kit, Biolegend, #423106), transferred into a 96-well plate and incubated for 20 min at 4 °C in the dark. Cells were washed two times with 150 µL PBS (centrifuged 5 min at 400 × g at 4 °C) and resuspended in 50 µL Fc-block (1:50, Purified Rat Anti-Mouse CD16/CD32, BD, #553142) diluted in FACS buffer (1x PBS/2% FBS/2mM, EDTA pH 8.0). Cells were incubated for 20 min at 4 °C in the dark and washed two times with 150 μl PBS (centrifuged 5 min at 400 × g at 4 °C). All cells were fixed for 30 min with BD Cytofix/Cytoperm solution (BD, #554722) and stored in PBS overnight. For the extracellular and intracellular fluorescent cell staining, cells were permeabilized with BD Perm/Wash buffer (BD, #554723) for 15 min. T lymphocytes were evaluated using the following antibodies: rat anti-mouse IL-17A (TC11-18H10.1, 1/50; Biolegend, #506922), rat anti-mouse RORγt (AFKJS-9, 1/44, eBiosciences, #17-6988-82), rat anti-mouse CD3 (17A2, 1/88, Biolegend, #100241), rat anti-mouse CD45 (30-F11, 1/88, BD, #564225), rat anti-mouse CD4 (RM4-5, 1/700, Biolegend, #100548), rat anti-mouse IFN-γ (XMG1.2, 1/175, eBiosciences, #61-7311-82), rat anti-mouse FOXP3 (FJK-16s, 1/200; ThermoFisher, #48-5773-82), rat anti-mouse CD8 (53-6.7, 1/700, Biolegend, #100710). Optimal staining concentrations of all antibodies were evaluated before staining. Cells were incubated with FACS buffer diluted antibodies for 30 min at 4 °C in the dark. Samples were washed twice with 150 μL of BD Perm/Wash buffer, resuspended in 200 µL PBS and acquired using Quanteon NovoCyte (NovoCyte Quanteon 4025, Agilent). All acquired data were analyzed using FlowJo™ Software (version 10.7.2, BD, 2019). Fluorescence minus one controls (FMOs) were used for each antibody-fluorophore combination to properly evaluate signal-positive and -negative cells. Single antibody-stained UltraComp eBeads™ Compensation Beads (Fisher Scientific, Ref: 01-2222-42) were used to create the compensation matrix in FlowJo. Compensation samples were gated on the population of compensation beads within the FSC and SSC and the positive and negative population for the corresponding antibody were identified. After applying the compensation matrix, samples underwent the following gating strategy: (1) identifying single live cells in a FSC-A vs. FSC-H plot; (2) selecting live cells in a Zombie NIR vs. SSC plot; (3) identifying single, live CD45^+^ cells in a CD45^+^ vs. SSC plot. All samples with less than 1000 events in this plot were removed from the analysis. FACS analysis of isolated lymphocytes was performed blinded and gating strategy was verified by two persons.

### Illumina 16S rRNA gene sequencing and analysis

Bacterial DNA extraction from colonic and ileal content was performed as described previously (*16*). A Qubit® dsDNA HS assay kit was used to quantify sample inputs. The V4 region of the 16S rRNA gene was amplified using dual-index primers described by Kozich et al (*54*). Library preparation was performed according to manufacturer’s protocol using the Quick-16S™ NGS Library Prep Kit (Zymo Research, Irvine, CA, USA, #D6400). The pooled libraries were sequenced on an Illumina MiSeq using MiSeq®Reagent Kit v2 (500-cycle) (Illumina, #MS_102_2003). All raw sequencing data have been uploaded to the European Nucleotide Archive (ENA) at EMBL-EBI under the study accession number PRJEB60278. The program mothur (v1.44.3) (*55*) was used to process the reads according to the MiSeq SOP (*54*). For gnotobiotic samples, taxonomy was assigned using a k-nearest neighbor consensus approach against a custom reference database corresponding to the SM14 taxa and potential contaminants (*Citrobacter rodentium*, *Lactococcus lactis* subsp. *cremoris*, *Staphylococcus aureus*, and *S. epidermis*). For SPF samples, taxonomy was assigned using the Wang approach against the SILVA v132 database. Count data was normalized by computing relative abundance.

### Construction of the phylogenetic tree of SM14 constituent strains

The phylogenetic tree was constructed based on 16S rRNA gene sequences and analyzed with Geneious Prime version 2021.2.2. Accession numbers of the sequences were: AY271254.1 (*A. muciniphila*), AB510697.1 (*B. caccae*), EU136682.1 (*B. ovatus*), HQ012026.1 (*B. thetaiotaomicron*), AB050110.1 (*B. uniformis*), AB370251.1 (*B. intestinihominis*), AB626630.1 (*E. rectale*), HM245954.1 (*C. symbiosum*), AJ505973.1 (*M. formatexigens*), AF192152.1 (*D. piger*), AB011816.1 (*C. aerofaciens*), AJ413954.1 (*F. prausnitzii*), AJ312385.1 (*R. intestinalis*). A neighbor-joining tree build model was created using global alignment with free end and gaps, 65% similarity index cost matrix, and a Tamura-Nei genetic distance model.

### Metabolomics analysis of cecal contents using capillary electrophoresis-time of flight mass spectrometry (CE-TOF/MS)

Cecal metabolites were extracted from about 10 mg of a freeze-dried sample by vigorous shaking with 500 μL of 100% MeOH supplemented with 20 μM methionine sulfone as well as 20 µM D-camphor-10-sulfonic acid, as per internal standards. Four 3 mm zirconia beads (BioSpec Products, Bartlesville, OK, USA) and about 100 mg of 0.1 mm zirconia/silica beads (BioSpec Products, Bartlesville, OK, USA) were added to this mix. Afterwards, samples were shaked vigorously for 5 minutes using a Shake Master NEO (BioSpec Products, Bartlesville, Oklahoma, U.S.A.). Next, 500 μL of chloroform and 200 μL of Milli-Q water were added, and samples were shaken vigorously for 5 minutes again and followed by centrifugation at 4600 × g for 30 min at 4°C. The resulting supernatant was transferred to a 5 kDa cut-off filter column (Ultrafree MC-PLHCC 250/pk) for metabolome analysis (Human Metabolome Technologies, Tsuruoka, Yamagata, Japan). The flow-through was dried under vacuum. Residue was dissolved in 50 μL of Milli-Q water containing reference compounds (200 μM 3-aminopyrrolidine and 200 µM trimesic acid). The levels of extracted metabolites were measured by CE-TOF/MS in both, positive and negative modes, using an Agilent 7100 capillary electrophoresis system (Agilent Technologies, Waldbronn, Germany) equipped with an Agilent 6230 TOF LC/MS system (Agilent Technologies, Santa Clara, CA, USA).

### Metabolite-of-interest screening pipeline

Given that EAE phenotypes were disconnected from the overall metabolome pattern, we looked for single metabolites which might explain observed differences in EAE disease course. Thus, we implemented a screening pipeline, comprising 20 independent analyses, to identify potential metabolites-of-interest that might explain differences in EAE outcomes. These analyses included evaluation of the contribution of each metabolite to the variance of the PC1 and PC2 axes in a mulidimensional reduction PCA plot (**Extended Data Fig. 3b**), correlation analyses (**Extended Data Fig. 3c**), as well as group-wise comparisons of metabolite concentrations (**Extended Data Fig. 3d**). By combining information obtained from these analyses, our goal was to shortlist microbiota-induced cecal metabolites that enable prediction of either the overall disease course or the relapse occurrence in EAE-induced mice. We concluded that a potential metabolite-of-interest should fulfill the following five criteria (1 − 5): (1) an overall significantly different concentration between all 7 groups (**Extended Data Fig. 3c**, “SIG”; **Extended Data Fig. 3e** “SIG”), as determined by one-way ANOVA tests. As we have observed differences in EAE outcome on a group-based level, this should also be reflected in different concentrations of a metabolite-of-interest between the groups. (2) a non-significant contribution to the variance of the PC2 axis (**Fig. 3c**, “PC2”; **Extended Data Fig. 3b**, **Extended Data Fig. 3e** “PC2_POS”, “PC2_NEG”). As shown in **Fig. 3a**, different microbiota compositions were well reflected by the position of individual mice on the PC1 axis. Interestingly, differences in EAE status (EAE-induced *vs.* non-EAE-induced) were reflected by the position of individual mice on the PC2 axis. EAE-induced mice of all microbiota compositions generally provided lower values on the PC2 axis compared to their non-EAE-induced counterparts harboring the same microbiota composition. In summary, we made the following four observations (i – iv): (i) Different EAE phenotypes were a consequence of different microbiota compositions; (ii) Microbiota composition was well reflected by the position on the PC1 axis; (iii) EAE-induced mice of all microbiota compositions provided the same shift towards lower values on the PC2 axis compared to non-EAE-induced mice and (iv), as the shift towards lower values on the PC2 axis occurred in every microbiota composition, we concluded that these shifts occured independent from the EAE disease phenotype given the significant differences in EAE outcomes in mice harboring different microbiota compositions. In summary, we concluded that these shifts on the PC2 axis were either a direct result of EAE induction (independent from the disease phenotype) or a consequence of different microbiota colonization times since EAE-induced mice harbored the respective microbiota for 3 more weeks compared to non-EAE-induced mice of the same microbiota composition. Thus, we concluded that metabolites, which significantly contributed to the PC2 axis shift, were not causal to different EAE phenotypes. Instead, their concentrations in EAE-induced mice appeared to be a feedback effect from either longer colonization or EAE induction itself. In consequence, these metabolites were removed from the list of metabolites-of-interest. (3) a significant correlation with either the overall disease course (**Fig. 3c**, “AUC”; **Extended Data Fig. 3e**) or the mean score during the relapse phase (REL) in EAE-induced mice. (4) a significant correlation with the presence of *A. muciniphila* (**Fig. 3c**, “AM” **Extended Data Fig. 3e**) since we observed different EAE phenotypes based on presence or absence of *A. muciniphila.* (5) a significantly different concentration when comparing non-EAE-induced mice harboring microbiota compositions leading to different EAE phenotypes upon EAE induction (**Fig 3c, Extended Data Fig. 3d**). This criterion would allow for assessing the prediction aspect of cecal metabolite concentrations. As SM13-colonized and GF mice resulted in a moderate phenotype, SM01-colonized mice in an intermediate phenotype and SM14-colonized mice in a severe phenotype, we hypothesized that a potential metabolite of interest would reflect these differences by providing the following four properties (i − iv): (i) significantly different concentrations when comparing “SM13 – EAE” vs “SM14 – EAE” and “SM13 + EAE” vs “SM14 + EAE”; (ii) significantly different concentrations when comparing “SM13 – EAE” vs “SM01 – EAE” and “SM13 + EAE” vs “SM01 + EAE”; (iii) significantly different concentrations when comparing “SM01 – EAE” vs “SM14 – EAE” and “SM01 + EAE” vs “SM14 + EAE” and (iv) no significantly different concentrations when comparing “SM13 – EAE” vs “GF – EAE” and “SM13 + EAE” vs “GF + EAE”.

### Metatranscriptomics analysis

Flash-frozen cecal contents were stored at –80 °C until further processing. 1 mL RNAProtect Bacterial Reagent (Qiagen, #76506) was added to each sample and thawed on wet ice for 10 min. Samples were centrifuged at 10 000 × g, 4°C for 10 min and 250 µL acid-washed glass beads (212–300 μm), 500 µL of Buffer A (0.2 M NaCl, 0.2 M Trizma base, 20 mM EDTA pH 8), 210 µL of SDS 20% and 500 µL of phenol:chloroform (125:24:1) pH 4.3 were added to the pellet. Bead-beating on the highest frequency (30 Hz) was performed for 5 min using a mixer mill and the aqueous phase was recovered after centrifugation at 4°C for 3 min at 18 000 × g. 500 µL of phenol:chloroform (125:24:1) pH 4.3 was added to each sample and centrifuged as previously described. Again, the aqueous phase was recovered and 1/10 volume of 3M sodium acetate (pH ∼ 5.5) and 1 volume of ice-cold 100% ethanol was added and gently mixed by inversion. Samples were incubated for 20 min on wet ice and then washed twice with 500 µL of ice-cold 70% ethanol and centrifuged at 4°C for 20 min at 18 000 × g. Pellets were air-dried for 10 min and resuspended in 50 µL nuclease-free water. DNase treatment was performed by adding 10 µL 10X buffer, 40 µL nuclease-free water (to reach 100 µL final volume) and 2 µL DNase I (Thermo Scientific, DNase I, RNase-free kit, #EN0521) followed by 30 min incubation at 37°C. 1 µL EDTA 0.5M (per sample) was added and heat-inactivated for 10 min at 65°C. RNA purification was performed with the RNeasy Mini kit (QIAGEN, #74106) according to manufacturer instructions. RNA quantity and a quality were assessed using the RNA 6000 Nano Kit on an Agilent 2100 Bioanalyzer. Library preparation was performed using an Illumina® Stranded Total RNA Prep, Ligation with Ribo-Zero Plus kit (Illumina, #20040529). Pooled libraries were then run on a NextSeq 550 High Output flow cell 2×75bp up to 800M reads followed by NextSeq 550 Medium Output flow cell 2×75bp up to 260M reads. RNA sequencing files were pre-processed using kneaddata (https://github.com/biobakery/kneaddata). Adapters were removed using Trimmomatic (16) and fragments below 50% of the total expected read length (75 bp) were filtered out. BowTie2 (17) was used to map and remove contaminant reads corresponding to either rRNA databases or the *Mus musculus* genome. Clean fastq files were concatenated before passing to HUMAnN3 (18). A custom taxonomy table based on pooled 16S rRNA sequencing abundances was provided to metaPhlan for metagenome mapping. Unaligned reads were translated for protein identification using the UniRef90 database provided within HUMAnN3. Data for all samples were joined into a single table and normalized using count per million (CPM) method. Results were grouped by annotated protein product per individual. In case no annotation from UniRef90 transcript IDs was possible, distinct IDs were treated as separate gene products. Only gene products that provided >50 CPM in at least two of the eight investigated samples were included into downstream analyses. This resulted in 2213 transcripts being included into downstream analyses, representing 80 % to 85 % of the total CPM with no significant differences between the analyzed groups. CPM were recalculated to account for removed transcripts, followed by further analysis using the *edgeR* package (version 3.38.4) in R Studio (version 4.2.1). Multidimensional reduction of the transcriptome profiles was calculated using the *logFC* method within the *plotMDS.DGEList* function. Groupwise comparison of gene expression was calculated using the *exactTest()* function.

### Bacterial IgA coating index

Fecal samples stored at –20 °C were resuspended in 500 µL ice-cold sterile PBS per fecal pellet and mechanically homogenized using a plastic incolulation loop. Pellets were then thoroughly shaken for 20 min at 1100 rpm and 4 °C. After adding 2 × volume of ice-cold PBS, samples were centrifuged for 3 min at 100 × g and 4°C to sediment undissolved debris. Clear supernatant was recovered and passed through a 70 µm sieve (Imtec, #U3_10070_70) followed by centrifugation for 5 min at 10 000 × g and 4 °C to sediment bacteria. Supernatant was removed and pellet resuspended in 1 mL ice-cold PBS. Next, optical density of this suspension at 600 nm (OD_600_) was detected and approximate concentration of bacteria was estimated based on the assumption that OD_600_ = 1 equals 5 × 10^8^ bacteria per mL. The respective volume corresponding to 10^9^ bacteria was centrifuged for 5 min at 10 000 × g and 4 °C. Pellet was then resuspended in 400 µL 5 % goat serum (Gibco, #11540526) in PBS and incubated for 20 min on ice. After incubation, pellet was washed once in ice-cold PBS and centrifuged for 5 min at 10 000 × g and 4 °C. Pellet was then resuspended in 100 µL ice-cold PBS containing 4 µg of FITC-coupled goat anti-mouse IgA antibody (SouthernBiotech, Imtec Diagnostic, #1040-02). The ratio of 4 µg of the respective antibody to stain 10^9^ bacteria was previously evaluated to be the maximum amount of antibody that can be used without resulting in unspecific staining of non-IgA coated bacteria by using fecal samples from *Rag1*^−/−^ mice as non IgA-coated negative controls. Samples were then incubated for 30 min at 4 °C on a shaker at 800 rpm. After incubation, 1 mL ice-cold PBS was added followed by centrifugation for 5 min at 10 000 × g and 4 °C. Samples were then washed once in ice-cold PBS, either subjected for flow-cytometry detection or for separation of IgA^+^ and IgA^−^ bacteria. For immediate flow cytometric detection, pellets were resuspended in 200 µL DNA staining solution (0.9% NaCl in 0.1M HEPES, pH 7.2, 1.25 µM Invitrogen SYTO™ 60 Red Fluorescent Nucleic Acid Stain, Fisher Scientific, #10194852) followed by incubation for 20 min on ice. After washing once with PBS, pellets were resuspended in 100 µL PBS and run immediately on a Quanteon NovoCyte (NovoCyte Quanteon 4025, Agilent). For separation of IgA^+^ from IgA^−^ negative bacteria, we used the MACS cell separation system from Miltenyi. Pellets were resuspended in 100 µL staining buffer (5% goat serum in PBS) containing 10 µL anti-FITC microbeads (Miltenyi, #130-048-701) per 10^9^ bacteria, mixed well and incubaterd for 15 min at 4 °C. After end of incubation time, 1 mL of staining buffer was added, followed by centrifugation for 5 min at 5 000 × g. Pellet was then resuspended in 5 mL staining buffer per 10^9^ bacteria and loaded onto MACS LD separation columns (Miltenyi, #130-042-901) and flow-through containing the IgA^−^ fraction was collected. After removing columns from the magnet, the IgA^+^ fraction was flushed out and collected. The IgA^+^ fraction was then loaded on a MACS LS separation column (Miltenyi, #130-042-401). Flowthrough was collected and combined with the previous IgA^−^ fraction. After removing columns from the magnet, the IgA^+^ bacteria fraction was flushed out, collected and combined with the previous IgA^+^ fraction. Both combined fractions, IgA^+^ and IgA^−^ were then centrifuged for 10 min at 10’000 × g and 4 °C. Pellets were then resuspended in 1 mL PBS and subjected to two different downstream analyses: (1) To test purity of both fractions for each sample by flow cytometry, 10% of the suspension volume was used for bacterial DNA staining using SYTO™ 60 Red Fluorescent Nucleic Acid Stain as described above, whereas purity was generally >90% for both fractions; (2) To purify bacterial DNA for subsequent 16S rRNA gene sequencing of bacteria within the different fractions, 90% of the suspension volume was centrifuged for 10 min at 10 000 × g and 4 °C, supernatant was discarded and the dry pellet was stored at −20 °C. DNA isolation and 16S rRNA gene sequencing was then performed as described above. The IgA-coating index (*ICI*) for a given species *x* (*ICI_x_*) was calculated by the following equation, with *A_x_^+^* representing the strain-specific relative abundance in the IgA^+^ fraction and *A_x_^−^* representing the strain-specific relative abundance in the IgA^−^ negative fraction: 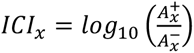. Strains were classified as “highly coated”, when 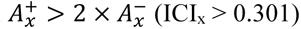 and as “low coated”, when ICI_x_ < −0.301.

### Soluble IgA measurements in fecal contents

We used 10 ng per well rabbit anti-mouse IgA (Novus Biologicals, #NB7506) for overnight coating of high binding 384 well plates (Greiner, #781061) in 20 µl per well of carbonate-bicarbonate buffer (Sigma, #C3041). Plates were then washed four times in wash buffer (10 mM Trizma Base, 154 mM NaCl, 1% (v/v) Tween-10). Next, 75 µL of a blocking buffer (15 mM Trizma acetate, 136 mM NaCl, 2 mM KCl, 1% (w/v) bovine serum albumin (BSA)) was added to each well and incubated for 2 h at RT. Following another washing step with was buffer, samples and standards were diluted in a dilution buffer (blocking buffer + 0.1% (w/v) Tween-20). As standards, we used a mouse IgA isotype control (Southern Biotech, #0106-01). 20 µL of the dilutions were added to each well and incubated for 90 min at RT. Following another washing step, a secondary goat anti-mouse IgA antibody, conjugated with alkaline phosphatase (Southern Biotech, # 1040-04), diluted 1:1000 in dilution buffer, was added. Secondary antibody was incubated at RT for 90 min and plates were washed four times. As a substrate, 1 phosphate tablet (Sigma, #S0642-200 TAB) was solubilized in 1 mM 1 mM 2-Amino-2-methyle-1-propanole and 0.1 mM MgCl2.6H2O. 40 µL of this substrate solution was added to each well, followed by incubation at 37 °C for 60 min. Final absorbance at 405 nm was detected using a SpectraMax Plus 384 Microplate Reader.

### Quantification of lipopolysaccharides levels in blood serum

To quantify lipopolysaccharide levels in blood serum from mice, full blood was incubated for 30 min at 37 °C, followed by centrifugation for 30 min at 3000 × rpm and RT. Serum supernatant was then stored at −80 °C until use. Quantification was performed using the Pierce Chromogenic Endotoxin Quantification Kit (ThermoScientific, #A39552) according to manufacturer’s instructions. Thawed serum samples were heat-shocked for 15 min at 70 °C and diluted 1:50 prior to performing the assay. After blank reduction, final endotoxin levels were calculated based on detected optical densities (OD) of supplied standards and using RStudio (version 4.2.1) using a 4-parameter nonlinear regression of standard ODs with the help of the *drc* package (version 3.0.1) applying the function *drm(OD∼concentration, fct=LL.4())*. Sample concentrations were then extracted using the *ED(type=”absolute”)* function of the same package.

### ELISA-based quantification of zonulin and occludin concentrations in blood serum

To measure concentrations of zonulin (ZO-1) and occludin (OCLN) in blood serum, we used the Mouse Tight junction protein ZO-1 ELISA Kit (MyBioSource, #MBS2603798) and the Mouse Occludin (OCLN) ELISA Kit (ReddotBiotech, #RD-OCLN-Mu), respectively, according to the manufacturer’s instructions. After blank reduction, final ZO-1 and OCLN concentrations were calculated based on detected optical densities (OD) of supplied standards and using R Studio (version 4.2.1) applying a 4-parameter nonlinear regression of standard ODs with the help of the *drc* package (version 3.0.1) using the function *drm(OD∼concentration, fct=LL.4())*. Sample concentrations were then extracted using the *ED(.., type=”absolute”)* function of the same package.

### Lipocalin-2 ELISA

To measure fecal lipocalin-2 levels, the fecal pellet were homogenized in 500 μl ice-cold PBS with 1% Tween 20. Samples were then subjected to, agitation for 20 min at 4 °C at 2000 rpm on a thermomixer, followed by centrifugation for 10 min at 21000 × g and 4 °C. Pellets were discarded, supernatants were harvested and stored at –20°C until further use. Final Lipocalin-2 detection was conducted using the Mouse Lipocalin-2/NGAL DuoSet Elisa, R&D Systems (# DY1857) according to the manufacturer’s instructions.

### Detection of bacterial relative abundances using quantitative real-time PCR

To detect relative abundances of commensal bacteria from fecal samples obtained from mice harboring reduced communities (SM01 to SM14) in a gnotobiotic setting, we followed a protocol published elsewhere (*21*) without modifications. Primer sequences for strain-specific quantification are listed in **Extended Data Table 2**.

### Detection of glycan-degrading enzyme activities from mouse fecal samples

To detect activities of α-fucosidase, α-galactosidase, β-glucosidase, β-*N*-acetyl-glucosaminidase and sulfatase from fecal samples stored at −20 °C, we followed the protocol published in detail elsewhere (*56*) without modifications.

### Data analysis

All figures and analyses were performed using RStudio (versions 4.1.3 and 4.2.1). For details see analysis-specific informations in this section.

### Illumina 16S rRNA gene sequencing-based analysis of complex microbial communities

Groupwise analysis of annotated reads was performed using RStudio (version 4.2.1) with an initial seed set at 8765. All operational taxonomic units (OTUs) not constituting at least 0.1% of reads within at least one group (group means) were removed from the analysis. Diversity indices were determined using the *diversity()* function of the *vegan* package (version 2.6.2). Nonmetric multidimensional scaling for Bray-Curtis distance matrices were calculated using the *metaMDS()* function of the *vegan* package and principal coordinate decomposition of weighted UniFrac distance matrices were calculated the *pcoa()* function of the *ape* package (version 5.6.2). All analyses were performed on OTU level, genus level and family level. OTUs and genera contributing most to community differences between selected groups were extracted using the *simper()* function of the *vegan* package.

## Acknowledgements

This work was supported by the following grants in the laboratory of M.S.D.: Luxembourg National Research Fund (FNR) CORE grants (C15/BM/10318186 and C18/BM/12585940) and BRIDGES grant (22/17426243) to M. S. D; and FNR INTER Mobility grant (16/11455695) to M. S. D.; FNR AFR bilateral grant (15/11228353) support to M.N.; FNR AFR individual PhD fellowship (11602973) to A. P.; E.T.G. was supported by the FNR PRIDE grant (PRIDE17/11823097) and the Fondation du Pélican de Mie et Pierre Hippert-Faber, under the aegis of the Fondation de Luxembourg. We thank MEDICE Arzneimittel Pütter GmbH & Co. KG, Germany and Theralution GmbH, Germany for funding through the public–private partnership FNR BRIDGES grant (22/17426243). E.M. is supported by The Japan Society for the Promotion of Science (JSPS) KAKENHI (19K05907), Lotte Foundation, and Astellas Foundation for Research on Metabolic Disorders. S.F. (including CE-TOF/MS metabolomic analyses) was supported by JSPS KAKENHI (22H03541), JST ERATO (JPMJER1902), AMED-CREST (JP22gm1010009), and the Food Science Institute Foundation. We acknowledge the National Cytometry Platform (NCP) for their assistance with flow cytometry. The NCP is supported by Luxembourg’s Ministry of Higher Education and Research (MESR) funding. We also thank the support of Christian Jäger, Xiangyi Dong and Floriane Gavotto from the Luxembourg Centre for Systems Biomedicine (LCSB), Metabolomics Platform for GC-MS analyses. We thank Kathy McCoy for providing us with *Muc2*^−/−^ mice.

## Data availability statement

The raw fastq files for this study from 16S rRNA gene sequencing and RNA sequencing have been deposited in the European Nucleotide Archive (ENA) at EMBL-EBI under accession number PRJEB60278 (https://www.ebi.ac.uk/ena/browser/view/PRJEB60278).

## Extended Data Figures

**Extended Data Figure 1:**
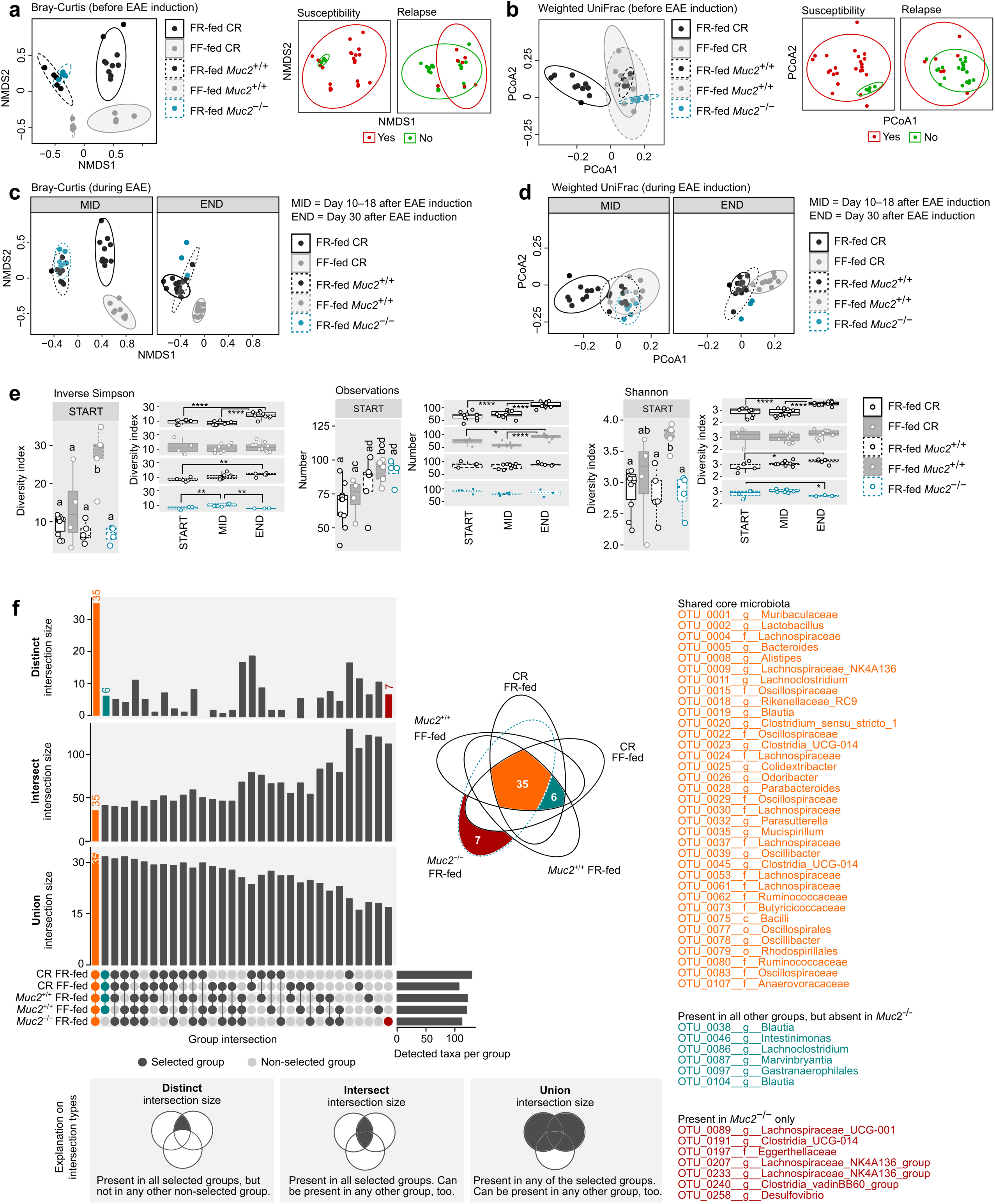
OTU level-based microbiota analysis of SPF-housed mice. 16S rRNA gene-based Illumina sequencing data analyses of DNA isolated from fecal samples. All samples were analyzed together using the same analysis pipeline. Only those OTUs that provided a mean relative abundance of at least 0.01% in at least one group were included in the analyses. **a, b)** Multifactor dimensionality reduction (**a**, NMDS plot based on a Bray-Curtis distance matrix; **b**, Principal coordinate analysis (PCoA) using a weighted UniFrac distance matrix) of microbiota composition based on OTUs. Fecal samples collected the day before EAE induction. **a**) Left panel shows clustering of samples by group, middle and right panel depict the same samples on the same two-dimensional scale as the left panel, colored by susceptibility to EAE or occurrence of relapse in the same mouse from which the respective sample was obtained. Ellipses represent 95% confidence interval for each group. The minor differences between groups in b), compared to more prominent differences between groups in a) indicate that the detected differences in the NMDS plot (a) are mostly based on phylogenetically related OTUs. **c, d)** Multifactor dimensionality reduction (c, NMDS plot based on a Bray-Curtis distance matrix; d, Principal coordinate analysis (PCoA) using a weighted UniFrac distance matrix) of microbiota composition based on OTUs. Fecal samples were collected at day 10 to 18 (“MID”) or at day 30 (“END”) after EAE induction. Ellipses represent 95% confidence interval for each group. Detected shifts during EAE disease course using both methods of multifactor dimensionality reduction were disconnected from group EAE phenotype (Fig. 1). **e)** Alpha-diversity analysis of microbiota composition before and during EAE, separated by groups and diversity indices. Fecal samples collected the day before EAE induction (“START”), at day 10 to 18 (“MID”) or at day 30 (“END”) after EAE induction. Observations = number of detected taxa (OTUs). Statistics: statistical differences were calculated by One-way ANOVA per panel, followed by a Tukey’s post-hoc test for groupwise comparison. Statistical differences between groups before EAE induction (START) are indicated by a compact letter display; two groups sharing an assigned letter, p > 0.05; two groups not sharing an assigned letter, p < 0.05 (without further distinction). Statistical differences between different timepoints within the same groups are indicated by asterisks: *, p < 0.05; **, p < 0.01; ***, p < 0.001; ****, p < 0.0001. Non-significant comparisons are not shown. **f)** Intersection analysis to identify taxa, which are either present or absent only in *Muc2^−/−^*mice. Taxa were considered “present” within a certain group, when group mean relative abundance of a given taxon was > 0.01%. Otherwise, taxa were considered “absent” in the respective group. Left panels: intersection sizes of OTUs between the five groups. Three different intersection sizes are shown (“Distinct”, “Intersect”, and “Union”), which are explained in the lower panels. Middle panel: Venn diagram highlighting the number of shared and distinct OTUs between the groups. Dark red, taxa only present in *Muc2^−/−^* mice; teal, taxa not present in *Muc2^−/−^* mice, but in all other mice; orange, core microbiome shared by mice of all five groups. List of OTUs belonging to these three groups provided on the right.

**Extended Data Figure 2:**
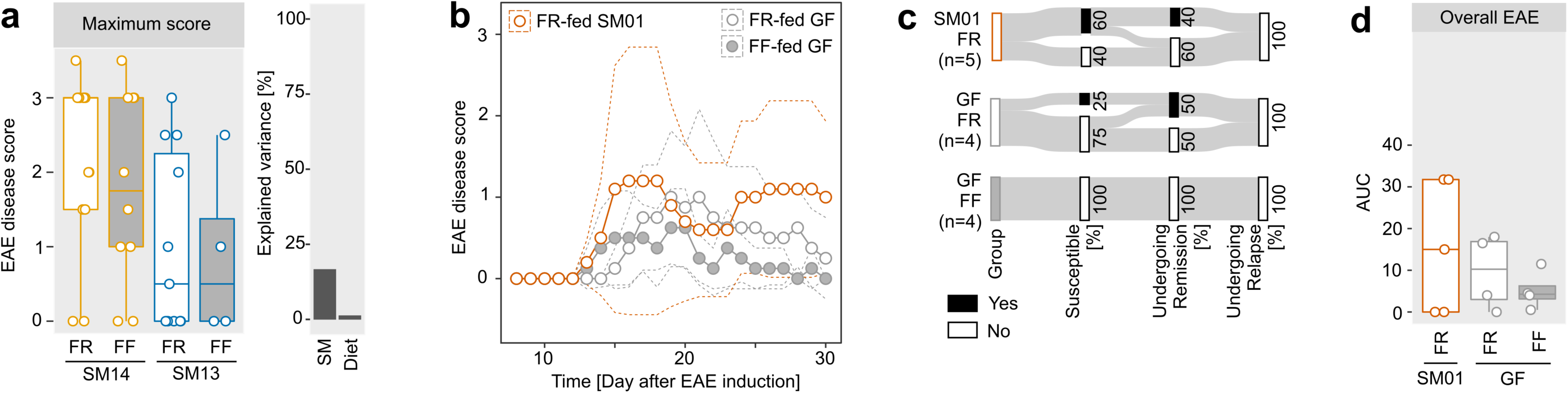
EAE disease course in germ-free as well as SM01-, SM13- and SM14-colonized mice. **a)** Left panel: maximum achieved EAE score per individual within FR-fed SM13-colonized mice (n=11), FF-fed SM13-colonized mice (n=4), FR-fed SM14-colonized mice (n=13) and FF-fed SM14-colonized mice (n=10). Each mouse depicted by a separate dot in the boxplot. Statistics: One-way ANOVA followed by Tukey’s post-hoc test. No statistically significant group differences observed. Right panel: Percentage of variance explained by diet (“diet”, FR vs. FF) and SM combination (“SM”, SM13 vs. SM14) as determined by eta-squared (η^2^) calculation when comparing the maximum achieved EAE score during the 30 d disease observation period between FR- and FF-fed SM14- and SM13-colonized mice. **b–d)** GF C57BL/6 mice were either monocolonized with *A. muciniphila* (SM01) or remained germ free (GF). GF mice were fed either FR or FF diet. SM01 mice were only fed FR diet. EAE was induced in GF mice 20 d after diet switch. EAE was induced in SM01-colonized mice 25 d after initial colonization. Disease course in all groups was observed for 30 days after EAE induction. **b)** EAE disease scores as a function of time (days after EAE induction) for FR-fed GF mice (n=4), FF-fed GF mice (n=4) and FR-fed SM01-colonized mice (n=5). Dots represent daily group means. Dotted lines represent SD. **c)** Sankey diagram of key event occurrence (in % of all mice within one group) during EAE. Same event definition as in Fig. 1e. **d)** Area-under-the-curve (AUC) analysis of the disease course depicted in b). Each mouse depicted by a separate point. Statistics: One-way ANOVA followed by Tukey’s post-hoc test. No statistically significant group differences observed.

**Extended Data Figure 3:**
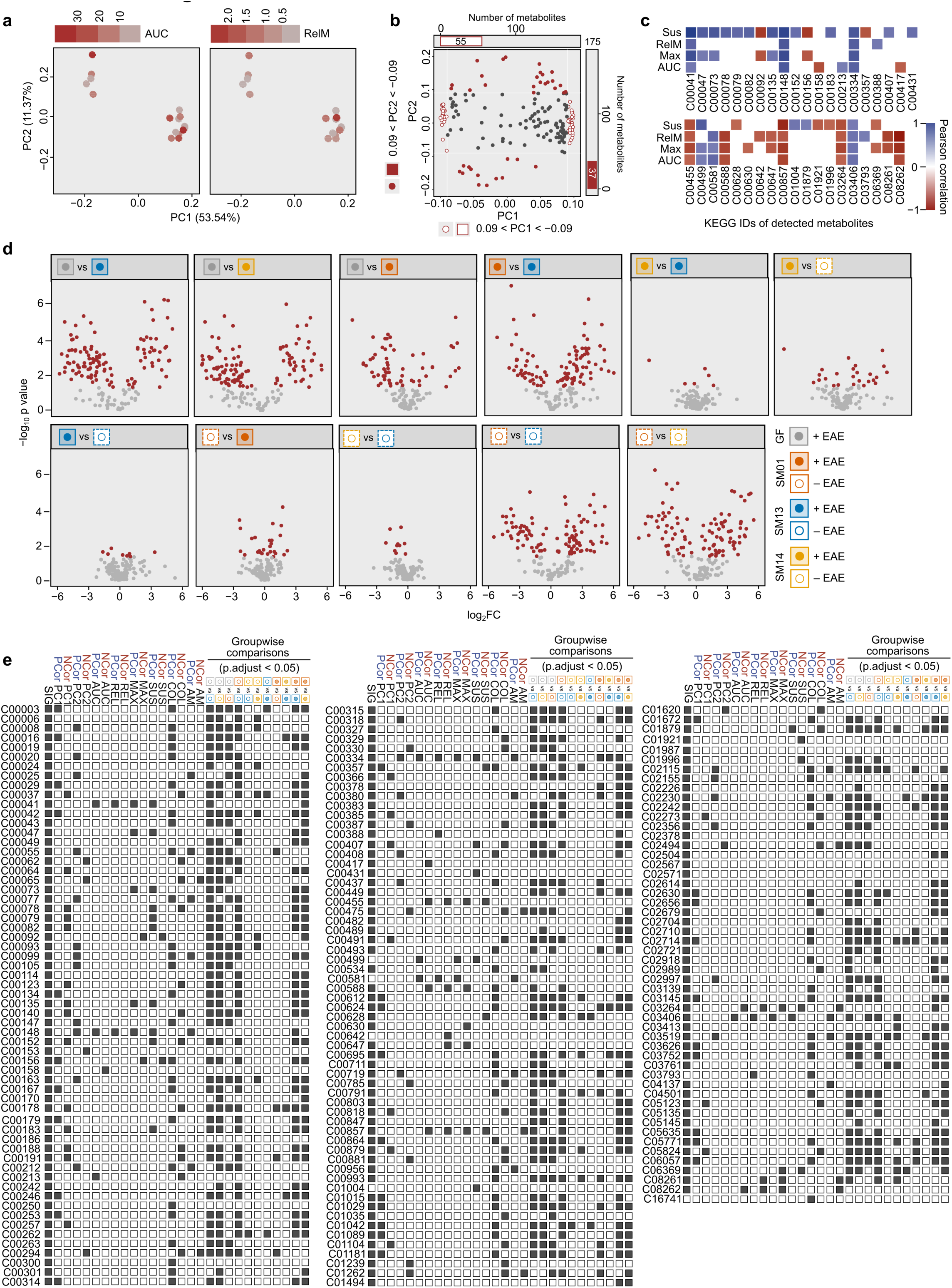
Metabolite-of-interest analysis pipeline. **a)** Principal component analysis (PCA) of log-normalized metabolite concentrations using a Euclidian distance matrix. Each dot represents an individual EAE-induced mouse, independent of microbiota composition. Dot colors represent EAE severity in each individual mouse (n=16), with strong EAE phenotypes colored in red. Severity represented by either AUC values of the individual EAE disease course as a function of time (left panel) or the mean score during the relapse phase (“RelM”; right panel). EAE phenotypes were disconnected from the overall metabolome pattern. **b)** Biplot showing the loadings of each metabolite, indicating significant positive or negative contribution to the PC1 and PC2 axes of a principal component analysis (PCA) of log-normalized metabolite concentrations using a Euclidian distance matrix (panel a, Fig. 3a). Each metabolite represented by one dot. Metabolites were classified as having a significant contribution when their loadings were either smaller than −0.09 or higher than +0.09 for both, PC1 (white circles with red lines) and PC2 (red circles). The total number of metabolites with significant contributions is shown on the top (for PC1) or the right (for PC2). **c)** Pearson’s correlation of metabolite concentrations with key EAE-associated readouts: “Sus” = Occurrence of susceptibility (categorical); “RelM” = Mean EAE disease score during relapse phase (numerical); “Max” = maximum achieved EAE score (numerical); “AUC” = Area-under-the-curve analysis of daily EAE scores as a function of time (numerical). Only samples from EAE-induced mice were used for analysis. Significant correlations are shown in either blue (positive) or red (negative). Non-significant correlations were removed from the figure. Metabolites are listed under their KEGG-ID. A list of metabolite names with corresponding KEGG-IDs is provided in Extended Data Table 3. **d)** Volcano plot of groupwise comparison for metabolite concentrations between the groups as indicated by the color-coded legend. Each dot represents one metabolite. Significantly different expressed metabolites are highlighted in red. Benjamini-Hochberg correction was used to adjust p-values. **e)** Criteria intersection analysis of all 175 detected metabolites. Metabolites are listed under their KEGG-ID. Fulfilled criteria indicated by a dark grey square; not fulfilled criteria indicated by a white square. Criteria were categorized into “correlation criteria”, summarizing the results of either statistically significant positive (“PCor”) or statistically significant negative Spearman correlation (“NCor”) across all samples of all groups (both, EAE-induced and non-induced mice) and “groupwise comparison criteria”. Correlations referring to EAE-associated criteria (“AUC”, Area under the curve of the disease course; “MAX”, maximum EAE score; “SUS”, susceptibility to EAE induction as defined in Fig 1c; “REL”, occurrence of relapse during the last 5 days of EAE course as defined in Fig 1e) were calculated using samples of EAE-induced mice only. Groupwise comparisons include metabolites found to be significantly different between two given groups based on an unpaired t-tests of log-normalized concentrations, using a BH-adjusted p-value (p.adjust) as the significance criterion (p.adjust < 0.05).

**Extended Data Figure 4:**
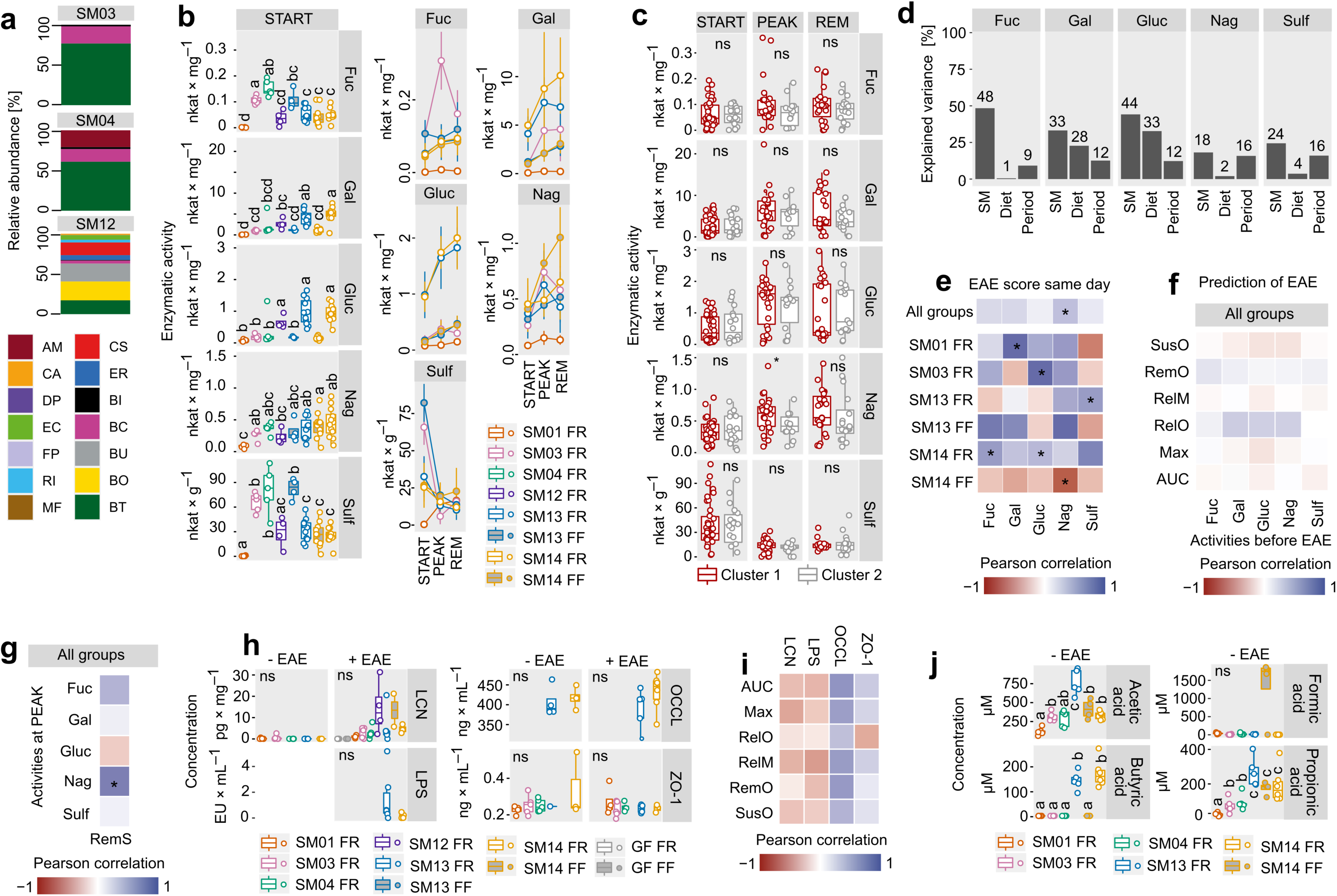
Analysis of barrier integrity and mucin-associated glycan degrading capacities of reduced microbiota compositions. **a)** Colonization verification. Relative abundance of bacterial strains in SM03-, SM04- and SM12-colonized mice after initial colonization as detected by qPCR using strain-specific primers (Extended Data Table 2). **b–g)** Given that two of the three SM combinations which resulted in a “severe” phenotype (Fig. 4c, Fig. 5c) comprised the full set of mucin glycan-degrading strains (*A. muciniphila, B. caccae, B. instinihominis, B. thetaiotaomicron*) within the SM14 community and given that increased activities of microbial mucin-glycan degrading enzymes, as detected from fecal samples, were previously reported to be associated with increased susceptiblity towards enteropathogenic infections (*16*), we detected activities of *N*-acetylglucosaminidase (Nag), α-fucosidase (Fuc) and sulfatase (Sulf) (*56*), which are involved in degradation of mucin-associated glycan structures (*16*). As controls, we also detected activities of enzymes involved in fermentation of mostly plant-derived fiber structures: α-glucosidase (Gluc) and β-galactosidase (Gal). Enzyme activities were measured at three different timepoints (“periods”) during the 30 d of EAE disease course: Before EAE induction (“START”), during the phase with the maximum EAE score (“PEAK”) which corresponded to day 14 to day 20, dependent on the individual, as well as during the remission phase (“REM”) which corresponded to day 21 to day 25 after EAE induction. **b)** Left panel: Boxplot of enzyme activities of Fuc, Gal, Gluc, Nag and Sulf as determined from fecal samples collected before EAE induction (START). Each dot represents one individual mouse. Statistics: statistical differences were calculated by One-way ANOVA per enzyme, followed by a Tukey’s post-hoc test for groupwise comparison. Statistical differences between groups before EAE induction (START) are indicated by a compact letter display: two groups sharing an assigned letter, p > 0.05; two groups not sharing an assigned letter, p < 0.05 (without further distinction). Right panel: Changes of enzyme activities per group during EAE disease course from START over PEAK to REM. Dots represent group means, vertical lines SD. Not all groups groups were detected at PEAK or REM. **c)** Boxplot of enzyme activities of Fuc, Gal, Gluc, Nag and Sulf as determined from fecal samples in EAE-induced mice, based on cluster affiliation and period after EAE induction. Same period definition as above; same cluster definition as in Fig. 5e. Each dot represents one individual mouse. Statistics: student’s t-test. *, p < 0.05; ns, non significant. **d)** Variance in enzymatic activities Fuc, Nag, Sulf, Gluc and Gal explained by microbiota composition (“SM”), diet (“Diet”) or period (“Period”; START vs. PEAK vs. REM) as determined by eta-squared (η^2^) calculation. While the microbiota composition turned out to be most determining factor for the different enzyme activities, diet only explained a considerable proportion of the variance for the two plant glycan-derived fiber-degrading enzymes Gluc and Gal. The timepoint of sampling during EAE course explained between 9% and 16% of the variances, suggesting that enzyme activities are, at least in part, either impacted by the current state of neuroinflammation or involved in mediating the current state of neuroinflammation **e)** Pearson’s correlation coefficients of fecal enzymatic activities of Fuc, Nag, Sulf, Gluc and Gal in EAE-induced mice with EAE scores at the same day of fecal sampling for each individual. Upper panel: all mice of all microbiota–diet combinations combined (“All groups”). Lower panel: only those mice were included in correlation analyses that belonged to the respective colonization–diet combinations. Statistically significant correlations (p < 0.05) are marked with * without further distinction. Only Nag activities correlated significantly with EAE scores at the same day when considering all mice independent from the microbiota composition. We observed considerable differences when including only mice harboring the same microbiota composition, or the same diet or both, into such analyses (lower panel), highlighting that enzyme activity-EAE readout correlations strongly depended on the microbiota composition. **f)** Pearson’s coefficients of correlations of enzymatic activities Fuc, Nag, Sulf, Gluc and Gal in fecal samples of EAE-induced mice before EAE induction with key EAE-associated readouts after EAE-induction in the same individuals. SusO, occurrence of susceptibility; RemO, occurrence of remission; RelM, mean EAE score during relapse phase; RelO, occurrence of disease relapse; Max, maximum achieved EAE score; AUC, area-under-the-curve analysis of EAE score as a function of time. All mice of all groups combined. No statistically significant correlations were determined. Consequently, no enzyme activity before EAE induction could predict any of the six key EAE-associated readouts when combining all mice of all colonization–diet combinations. **g)** Pearson’s coefficients of correlations of enzyme activities during PEAK with EAE scores during remission phase (“RemS”) of the same indiviual. Mice from all microbiota–diet combinations combined. Only Nag activites during the Peak phase could be used to predict EAE scores during the remission phase. Nag activities correlated positively with higher EAE scores, or in other words, less remission. This correlation was significant, but relatively weak. **h–j)** Given the significant correlation of Nag activities with certain EAE-associated readouts, we next addressed whether increased Nag activities might contribute to decreased mucosal barrier integrity and whether this might impact EAE development. It has previously been shown that increased Nag activities were associated with impairment of the intestinal mucus layer and increased lipocalin secretion into the intestinal lumen (*16*). Furthermore, reduced mucosal barrier integrity is discussed as a contributing factor to MS pathology (*57, 58*). Since we did not assess mucus layer thickness directly, we detected indirect measures of mucosal barrier integrity, such as serum levels of lipopolysaccharides (LPS), occludin and zonulin, as well as lipocalin concentrations from fecal samples. Additionally, we assessed short-chain fatty acid (SCFAs) concentrations given their contribution to maintenance of mucosal barrier integrity (*59, 60*) and the disease-alleviating properties of propionate in MS patients (*4*) and EAE-induced mice (*61*). **h)** Fecal concentrations of lipocalin (LCN) and serum concentrations of lipopolysaccharide (LPS), occludin (OCCL) and zonulin (ZO-1) as determined either from non-EAE-induced mice (−EAE) or in EAE-induced mice, 30 d after EAE-induction (+EAE). Some group/readout/timepoint combinations were not determined. Statistics: Statistical differences were calculated by One-way ANOVA per panel. ns, no statistically significant differences. **i)** Pearson’s coefficients of correlations between fecal LCN concentrations, serum OCCL concentrations, serum LPS concentrations and serum LCN concentrations before induction of EAE with key EAE-associated readouts in the same individual across all tested microbiota–diet combinations. SusO, occurrence of susceptibility; RemO, occurrence of remission; RelM, mean EAE score during relapse phase; RelO, occurrence of disease relapse; Max, maximum achieved EAE score; AUC, area-under-the-curve analysis of EAE score as a function of time. No statistically significant correlations could be determined. **j)** Fecal concentrations of short-chain fatty acids (SCFA) in non-EAE-induced mice of certain microbiota–diet combinations. Statistics: statistical differences were calculated by One-way ANOVA per SCFA, followed by a Tukey’s post-hoc test for groupwise comparison. Statistical differences between groups are indicated by a compact letter display: two groups sharing an assigned letter, p > 0.05; two groups not sharing an assigned letter, p < 0.05 (without further distinction). SCFA concentrations were unrelated to EAE disease phenotypes upon EAE induction.

**Extended Data Figure 5:**
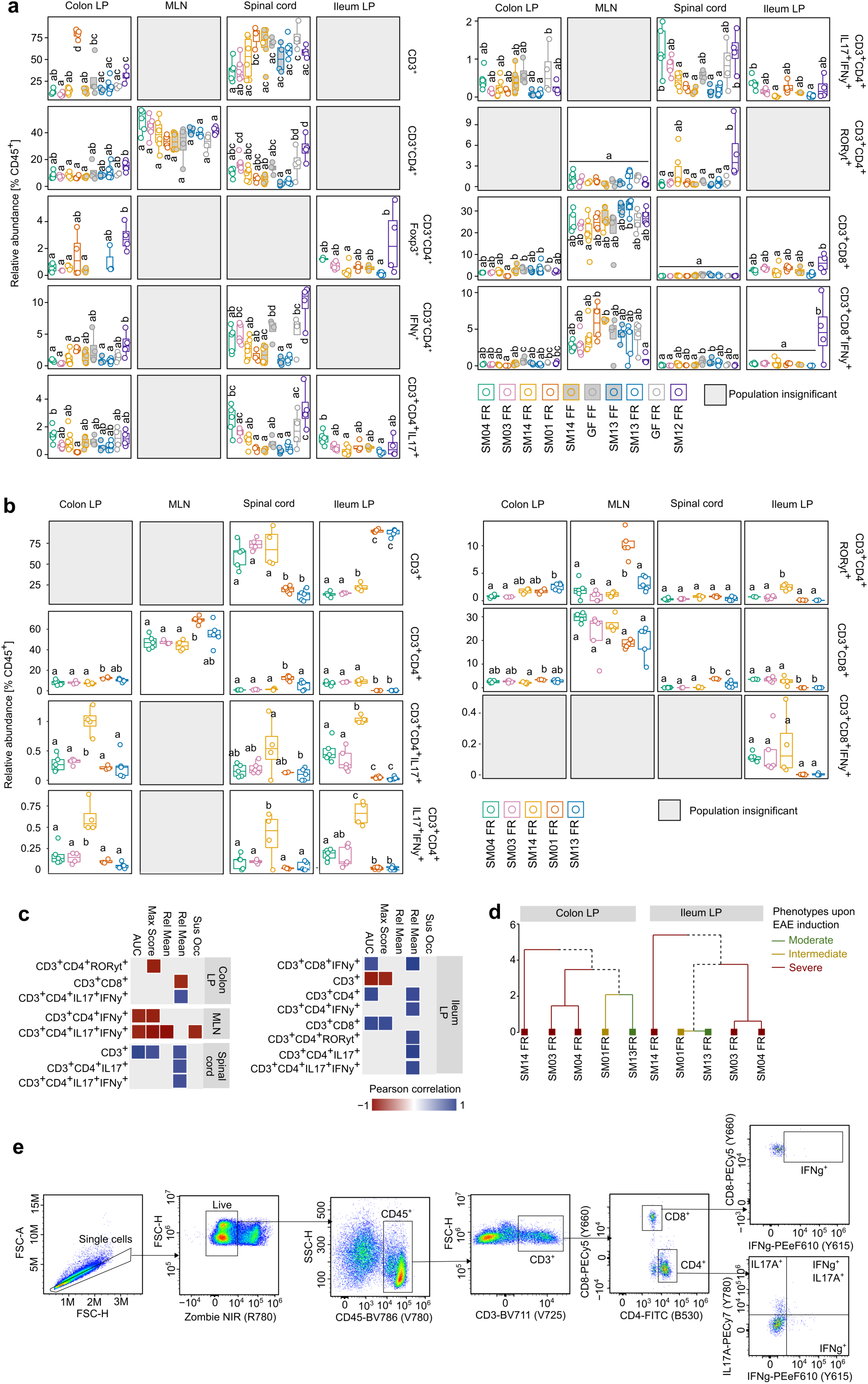
T cell polarisation in EAE-induced and non-EAE-induced mice. **a)** Boxplots of relative abundances of certain T cell populations in various organs. Group means of these populations are summarized in the heatmap shown in Fig. 5g. Colon, ileum, mesenteric lymphnodes and spinal cords were harvested followed by isolation of lymphocytes. Isolated lymphocates were subjected to flow cytometry-based analyses and relative abundances of subpopulations were calculated as percentage of CD45-expressing cells. LP, lamina propria. Each dot represents one individual. Groups are arranged based on their EAE group phenotype classification (Fig. 5c). Severe phenotype on the left: FR-fed SM04 (“SM04 FR”), FR-fed SM03 (“SM03 FR”), FR-fed SM14 (“SM14 FR”). Intermediate phenotype in the middle: FR-fed SM01 (“SM01 FR”), FF-fed SM14 (“SM14 FF”). Moderate phenotype on the right: FF-fed germ-free mice (“GF FF”), FF-fed SM13 (“SM13 FF”), FR-fed SM13 (“SM13 FR”), FR-fed germ-free (“GF FR”), FR-fed SM12 (“SM12 FRF”). Statistics: only population-organ combinations with statistically significant differences, as calculated by One-way ANOVA, are shown. Tukey’s honestly significant difference tests (Tukey HSD) were applied to subsequently determine statistically different value distribution between all groups within each population-organ combination. Statistical differences are indicated by a compact letter display. Two groups sharing an assigned letter, p > 0.05. Two groups not sharing an assigned letter, p < 0.05 (without further distinction). **b–d)** Analysis of T cell populations in non EAE-induced mice. Same populations were analyzed as shown in panel a. Only SM14-, SM13-, SM04-, SM03 and SM01-colonized mice were analyzed. All mice remained on a FR diet. Mice were analzyed after 25 days of colonization, representing the timepoint of EAE induction in EAE-induced mice. Colon, ileum, mesenteric lymphnodes and spinal cords were harvested followed by isolation of lymphocytes. Isolated lymphocytes were subjected to FACS-based analysis using the same antibody panels as used for EAE-induced mice. **b)** Boxplots of relative abundances of certain T cell populations in various organs. Each dot represents one invidual. Groups are arranged based on their EAE group phenotype classification (Fig. 5c). Statistics: only population-organ combinations with statistically significant differences are shown, as calculated by One-way ANOVA. Tukey’s honestly significant difference tests (Tukey HSD) were applied to subsequently determine statistically different value distribution between all groups within each population-organ combination. Statistical differences are indicated by a compact letter display. Two groups sharing an assigned letter, p > 0.05. Two groups not sharing an assigned letter, p < 0.05 (without further distinction). **c)** Pearson correlation matrix between group means of certain T cell subsets (vertical) isolated from different organs of non-EAE-induced mice with group means of EAE-associated readouts of EAE-induced mice harboring the same microbiota compositions (SM combinations, irrespective of the diet) as the non-EAE-induced mice. Non-significant correlations not shown. We found CD4^+^IL-17^+^ and CD4^+^IL-17^+^IFNγ^+^ cell populations in the colonic LP, spinal cords and the ileal LP to correlate significantly. This indicated that the microbiota composition already primed CD4^+^ T cells into a pro-inflammatory Th17 response before EAE-induction. **d)** Hierarchical group clustering of all 5 tested SM combinations based on scaled group means of significantly different T cells subsets, as determined by one-way ANOVA, separated by organ. Color codes indicate corresponding EAE group phenotype in EAE-induced mice, as determined in Fig. 5c. Population distribution in the colon that aligned best with emerging EAE group phenotypes upon EAE induction, indicating a crucial contribution of T cell priming in the colon by the microbiota to EAE disease course. **e)** Gating strategy to identify T cell subsets from colonic LP, MLN, SI and SC. Depicted is an example of a colonic LP-derived sample from a EAE-induced SM14-colonized mouse. Antibody-conjugated fluorochromes are indicated on the axis labels. Detection channels indicated in brackets. Black lines and squares indicate gates.

**Extended Data Figure 6:**
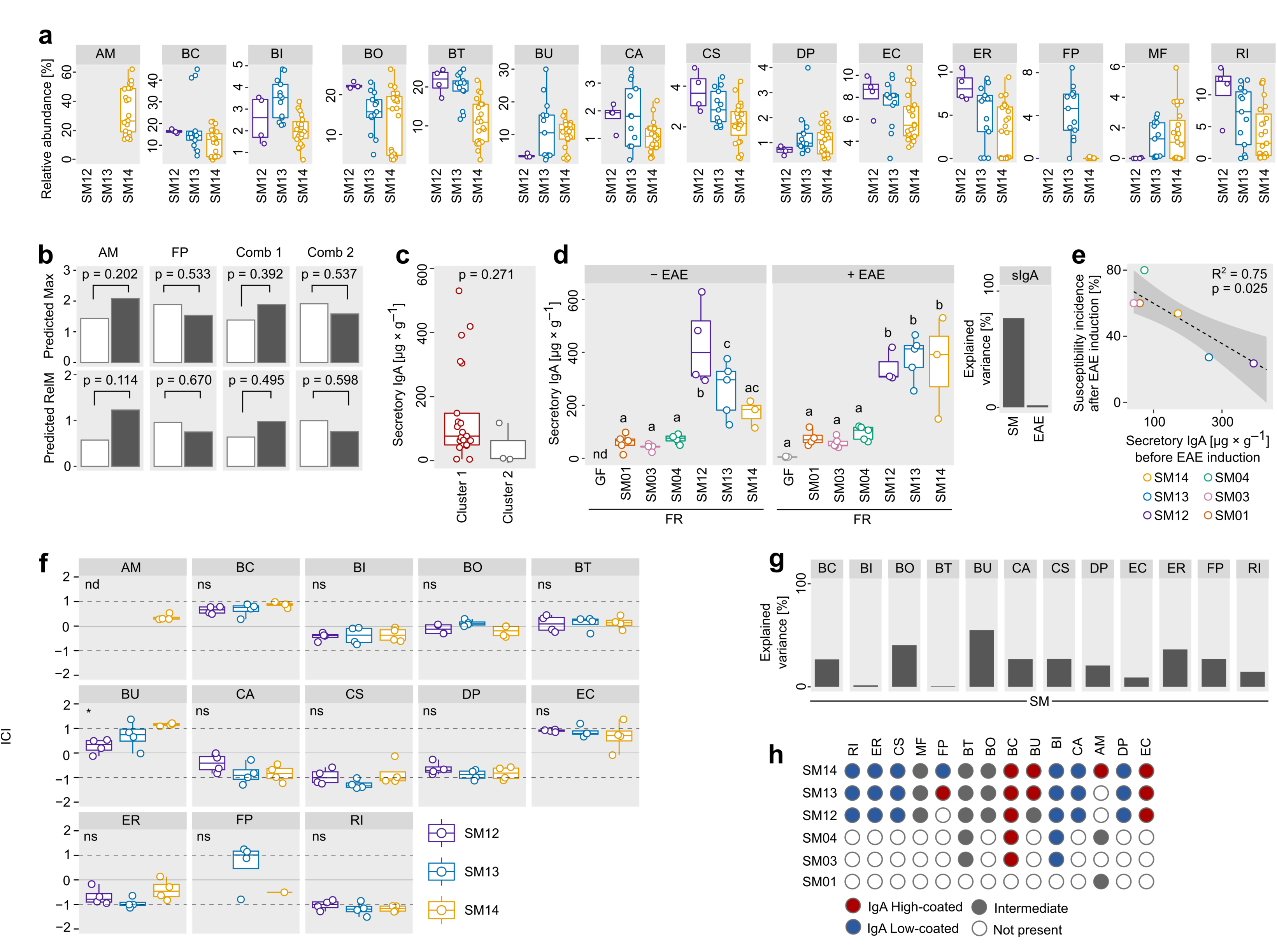
Microbiota-associated predictors for EAE development. **a)** Bacterial relative abundances of SM12-, SM13- and SM14-colonized mice, as detected by 16S rRNA gene sequencing, before induction of EAE. No distinction between diets. **b)** Linear mixed model regression for predicted maximum score during EAE (Max) and mean score during relapse phase (RelM) with presence of the strain as an independent variable and colonization as a random intercept effect. **c)** Concentrations of secretory IgA, determined from fecal samples and normalized to fecal weights. Fecal samples collected 30 d after EAE induction. Each dot represents one individual mouse. Mice grouped by individual EAE phenotype, as determined in Fig. 5e. Cluster 1, strong EAE symptoms; Cluster 2, mild EAE symptoms. Statistics: Wilcoxon rank-sum test. Intestinal secretory IgA levels after 30 d of EAE were independent (p = 0.271) from the respective individual EAE phenotype clusters. **d)** Concentrations of secretory IgA, determined from fecal samples and normalized to fecal weights. Fecal samples collected 30 d after EAE induction. Each dot represents one individual mouse. Mice grouped by colonization–diet combination and EAE induction status. “-EAE”: non-EAE-induced mice. “+ EAE”: EAE-induced mice. Statistics: Kruskal-Wallis test followed by Wilcoxon rank-sum test for groupwise comparisons. Statistical differences are indicated by a compact letter display. Two groups sharing an assigned letter, p > 0.05. Two groups not sharing an assigned letter, p < 0.05 (without further distinction). IgA levels after 30 d of EAE were significantly higher in mice harboring SM combinations with 12 or more strains. Right panel: variance of secretory IgA concentrations explained by either colonization (SM) or diet (Diet). **e)** Correlation between susceptibility (see Fig. 1e for definition) incidence in EAE-induced mice harboring a certain SM combination with mean secretory IgA levels in fecal samples obtained from non-EAE-induced mice harboring the same SM combination. Each dot represents one SM combination. Soluble IgA amounts, as detected from fecal samples, were a good predictor for susceptibility to disease upon EAE induction on a group level. **f)** IgA-coating index (ICI) of each strain dependent on SM combination. Each dot represents a sample obtained from one individual mouse. Statistics: One-way ANOVA. *, p < 0.05; ns, non significant. **g)** Variance in IgA-coating index (ICI) explained by background microbiota compositions (SM12, SM13 and SM14) **h)** Classification of SM14-constituent strains as IgA high-coated, low-coated or intermediate, dependend on microbiota composition. High coated, ICI > log(2); low coated, ICI < log(0.5); intermediate, log(0.5) < ICI < log(2).

**Extended Data Figure 7:**
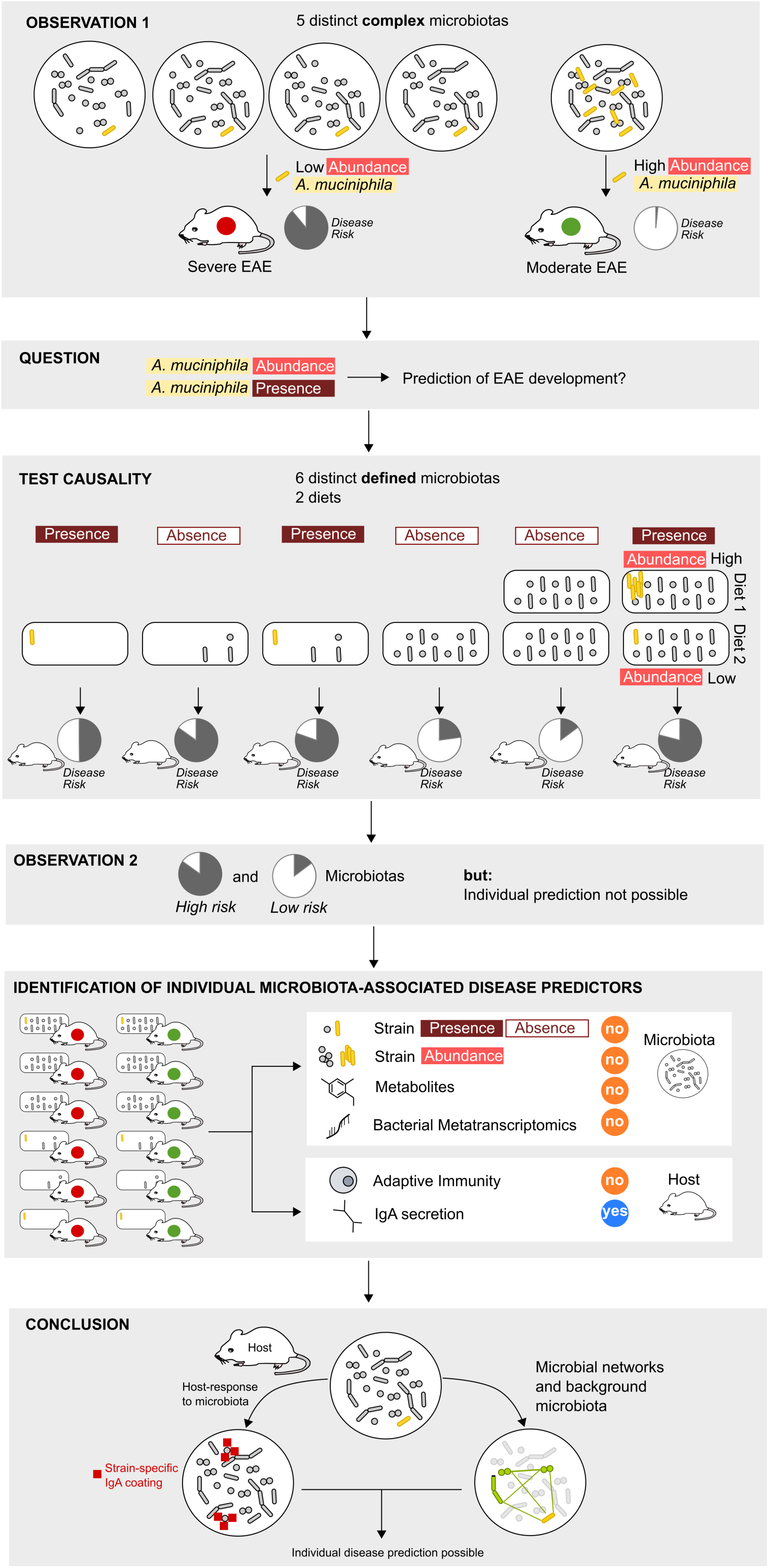
Graphical summary of experiments, analyses and conclusion. Based on the observation that low relative abundance of *Akkermansia muciniphila* (yellow rod-shape) in complex microbiota-harboring mice was associated with severe EAE development (red circle), and higher abundances of this species was associated with moderate disease (green circle) (OBSERVATION 1), we wondered whether either presence or the relative abundance of *A. muciniphila* before induction of EAE could predict subsequent disease development (QUESTION). To do so, we induced EAE in mice harboring 6 different combinations of a well-characterised 14-strain consortium under gnotobiotic conditions (TEST CAUSALITY). In mice harboring these microbiotas, *A. muciniphila* was either present or absent, and if present, *A. muciniphila* provided either high or low relative abundances. We found that distinct microbiota compositions resulted in “high risk” and “low risk” microbiota compositions, as determined by the proportion of mice developing severe disease (OBSERVATION 2). Given the fact that a certain microbiota composition did not result in equal disease susceptibility in every mouse harboring this particular microbiota, we aimed at identifying potential individual microbiota-associated disease predictors (IDENTIFICATION OF INDIVIDUAL MICROBIOTA-ASSOCIATED DISEASE PREDICTORS). We found that neither relative abundance, nor the presence or absence of *A. muciniphila*, or any other strain of the tested consortium, could reliably predict EAE development across distinct communities in individual mice. However, we identified the pre-EAE IgA coating index of another consortium member strain, *B. ovatus*, to significantly correlate with individual EAE outcome after disease induction, irrespective of the microbiota composition. Thus, we conclude that making predictions on EAE development based on microbiota characteristics (or in other words, assessing the individual disease risk) is possible. However, it needs to take into account inter-microbial interactions (“networks”) within a given, individual community as well as host-specific responses to a certain microbiota composition (CONCLUSION).

**Extended Data Figure 8:**
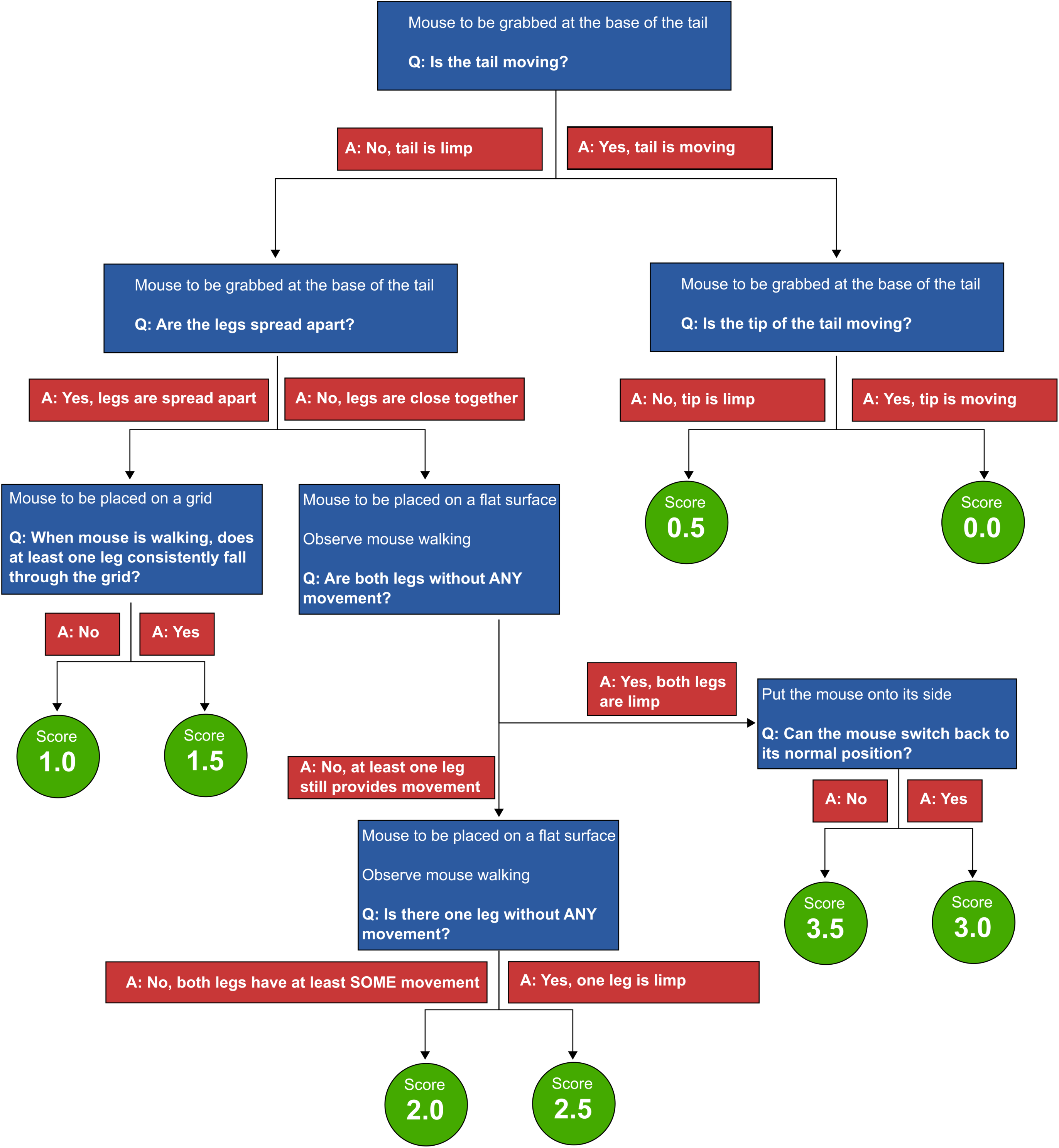
EAE scoring scheme. This decision tree depicts how daily assesment of EAE scores was performed. Blue boxes indicate instructions on mouse handling and EAE phenotype-associated questions (Q). Red boxes indicate possible answers (A) to the questions (Q). Green circles indicate the resulting EAE score. All arrows (answer options) are mutually exclusive.

